# Reconstitution of Alveolar Regeneration via novel DATPs by Inflammatory Niches

**DOI:** 10.1101/2020.06.14.151324

**Authors:** Jinwook Choi, Jong-Eun Park, Georgia Tsagkogeorga, Motoko Yanagita, Bon-Kyoung Koo, Namshik Han, Joo-Hyeon Lee

**Author notes:** Correspondence contact: Joo-Hyeon Lee.

## Abstract

Tissue regeneration involves a multi-step process composed of diverse cellular hierarchies and states that are also implicated in tissue dysfunction and pathogenesis. Here, we leveraged single-cell RNA sequencing analysis in combination with *in vivo* lineage tracing and organoid models to fine-map trajectories of alveolar lineage cells during injury repair and regeneration. We identified Damage-Associated Transient Progenitors (DATPs) as a distinct AT2-lineaged population arising during alveolar regeneration. Specifically, we found that IL-1β, secreted by interstitial macrophages, primes a subset of *Il1r1*^+^AT2 cells for conversion into DATPs, via a *Hif1a*-mediated glycolysis pathway, that are functional mediators for mature AT1 cell differentiation. Importantly, we show that chronic inflammation mediated by IL-1β prevents differentiation into AT1 cells, leading to aberrant accumulation of DATPs and impaired alveolar differentiation. Our step-wise fine-mapping of cell fate transitions demonstrates how the inflammatory niche impedes alveolar regeneration by directing stem cell fate behavior.

## Introduction

Maintenance of tissue homeostasis and repair following injury relies on the function of adult stem cells (Hogan et al., 2014; Li and Clevers, 2010; Wagers and Weissman, 2004). In the lung, barrier integrity of the epithelium is essential for protection against infection and efficient gas exchange. Lung tissue shows a slow cell turnover at steady state, but harbors regional-specific stem cells that quickly mobilize after tissue injury to replenish the epithelium (Hogan et al., 2014). In the alveoli, alveolar type 2 (AT2) cells maintain lung homeostasis and enable regeneration after injury by proliferating and differentiating into new alveolar type 1 (AT1) cells specialized for gas exchange (Adamson and Bowden, 1974; Barkauskas et al., 2013; Rock et al., 2011). Given the importance of AT2 cells, their self-renewal and differentiation must be tightly coordinated to maintain tissue integrity and efficient repair. Disruption of this balance can lead to life-threatening lung diseases (Hogan et al., 2014; Kotton and Morrisey, 2014). Recent studies have begun to suggest signaling pathways involved in the regulation of proliferation and differentiation of AT2 cells (Finn et al., 2019; Riemondy et al., 2019). However, it remains unclear which factors driven by injury trigger the activation of quiescent AT2 cells to differentiate towards AT1 cell fate and which differentiation trajectory they follow during lung regeneration.

Tissue repair is a complex process that involves dynamic crosstalk between stem cells and their respective niches. Physiological insults, such as a viral infection, are well known to instigate inflammation by triggering the activation or recruitment of immune cells to the affected tissue site (Medzhitov, 2008). In solid tissues, diverse immune cells of innate or adaptive immunity are even integral components of the niche, where they contribute to immune defence against infection and can sense environmental stimuli (Naik et al., 2018). Beyond the ability to clear pathogens, recent studies highlight how restoration of barrier integrity in epithelial organs such as the skin, gut, and lung after destruction is critically dependent on the immune system (Hsu et al., 2014; Klose and Artis, 2016; Lindemans et al., 2015; Naik et al., 2017). Lung epithelium is especially vulnerable to injury, as its surface is exposed to the external environment. In line with this, immune cells have been reported to be involved in lung homeostasis and restoration (Chen et al., 2012; Lechner et al., 2017; Westphalen et al., 2014). Recent advances have increased our insight into the critical role of paracrine niche-generated signals as key modulators of stem cell behaviors. In the distal lung, Pdgfra^+^ mesenchymal cells and vascular endothelial cells were identified as supportive niche cells (Barkauskas et al., 2013; Lee et al., 2014). More recently, mesenchymal cell subtypes including Wnt-responding and Wnt-producing fibroblasts were suggested to regulate stem cell properties and the cellular identity of AT2 cells (Lee et al., 2017; Nabhan et al., 2018; Zepp et al., 2017). However, our knowledge about the specific crosstalk between inflammatory cells and AT2 cells in regeneration remains limited. In particular, a fundamental question yet to be investigated is how the chronic inflammation impacts on tissue destruction, as it is likely caused by impaired stem cell function or regeneration process after injury, both processes that are poorly understood.

Here we set out to identify the lineage trajectory from AT2 toward AT1 cells during alveolar regeneration after injury. Single cell RNA-sequencing (scRNA-seq) analysis of *in vivo* AT2 lineage-labeled cells and *ex vivo* AT2 cell-derived organoids allowed us to delineate a precise differentiation trajectory in which AT2 cells adopt a ‘priming’ state followed by transition into ‘Damage-Associated Transition Progenitors (DATPs)’ prior to conversion into terminally differentiated AT1 cells. Importantly, we demonstrate that inflammatory niches driven by IL-1β and Hif1a signaling pathways orchestrate the regeneration process by triggering state-specific differentiation programs of AT2-lineage cells. Overall, our study reveals essential functions of inflammation in alveolar regeneration, providing new insights into how chronic inflammation impairs tissue restoration and leads to lung diseases.

## Results

### Reprograming of AT2 cells during alveolar regeneration after tissue injury

To define molecular identities and states of AT2 lineage cells responding to injury and undergoing regeneration, we treated AT2 reporter mice (*SPC-Cre^ERT2^;R26R^tdTomato^*) with tamoxifen, exposed them to PBS (control, homeostasis) or bleomycin (injury), and isolated lineage-labeled cells for scRNA-seq analysis at day 14 (acute injury) or 28 (resolution of injury) (Fig. 1A and Fig. S1A). Based on the expression of canonical AT1 and AT2 cell markers, we uncovered five distinct cell populations (Fig. 1B and Fig. S1B). Distribution of each cluster across time points allowed us to assess how AT2 cells changed during injury response and repair (Fig. 1C).

**Figure 1.**
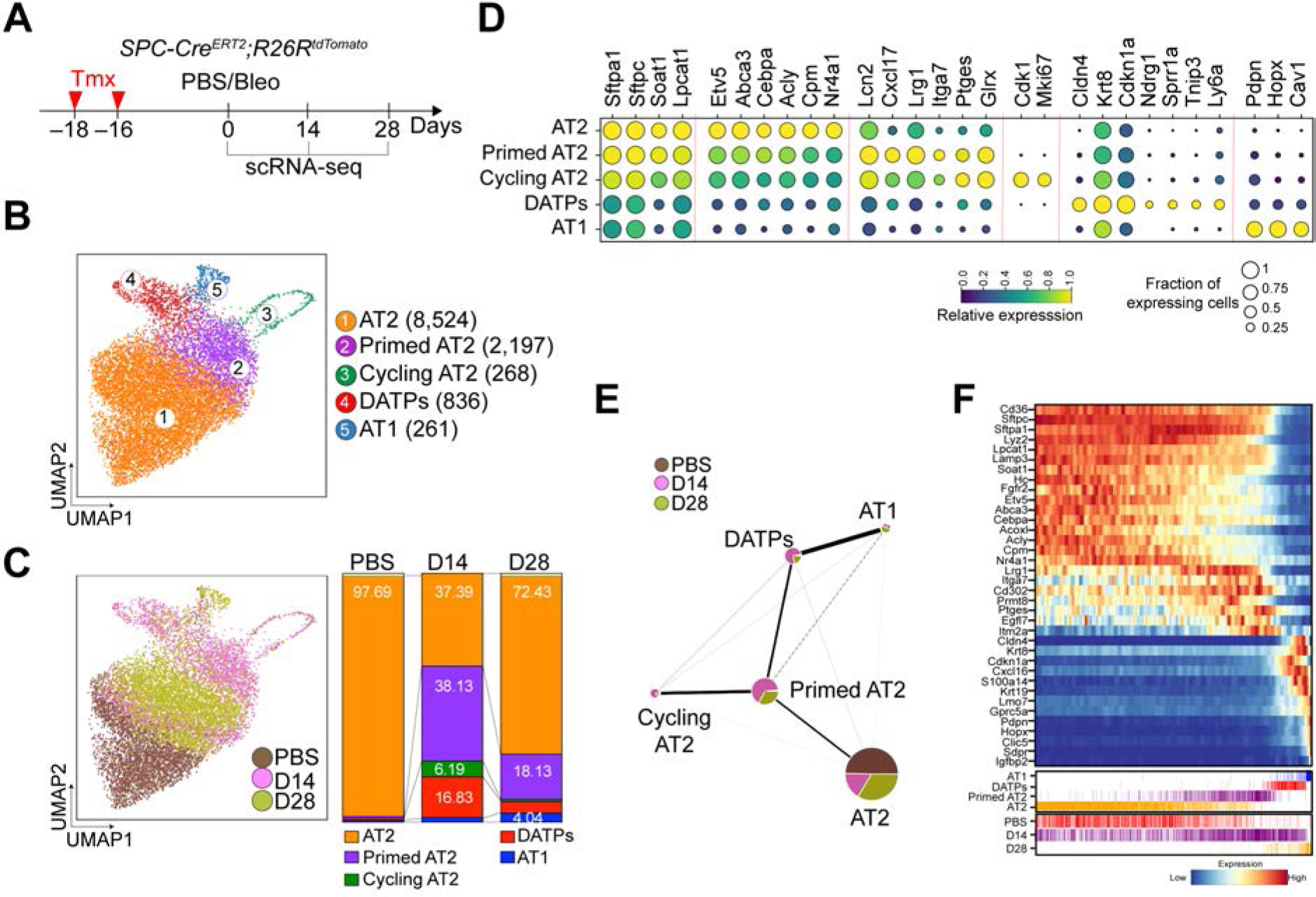
scRNA-seq reveals a dynamic lineage trajectory from AT2 cells to AT1 cells during alveolar regeneration after injury. **(A)** Schematics of experimental design for *SPC* lineage-labeled single cell isolation at indicated time points after bleomycin injury. **(B)** Clusters of *SPC* lineage-labeled alveolar cells (12,086) from 10xGenomics 3’ single-cell RNA sequencing (scRNA-seq) analysis visualized by UMAP, assigned by specific colors. Number of cells in the individual cluster is depicted in the figure. **(C)** Distribution of each cluster across indicated time points after injury. **(D)** Gene expression of key markers in each distinctive cluster. **(E)** Network topology among clusters from single cell data revealed by Partition-based graph abstraction (PAGA). Colors indicate the proportion of each cluster by time point. Each node in the PAGA graph represents a cluster and the weight of the lines represents the statistical measure of connectivity between clusters. **(F)** Heat map of gene expression profiles according to pseudotime trajectory. Lower color bars indicate cell types (upper panel) and actual time (bottom panel). **See also** Fig. S1.

As expected, lineage-labeled cells in uninjured mice, comprised mainly of AT2 cells (cluster 1) expressing canonical AT2 markers such as surfactant proteins (*Sftpc*, *Sftpa1*) (Fig.1, C and D). At day 14 post injury, three additional distinct populations had emerged while this AT2 cluster had become dramatically reduced (Fig. 1, C and D). Approximately 6% of lineage-labeled cells expressed proliferation/cell-cycle markers such as *Cdk1*, *Mki67*, and *Cenpa*, corresponding to Cycling AT2 cells (cAT2, cluster 3) (Fig. 1D and Fig. S1C). A second AT2-like state was highly prominent at this stage (cluster 2). This cluster showed similar expression levels of canonical AT2 markers including *Sftpc* but lower expression of genes that are involved in the lipid metabolism shown in homeostatic AT2 cells (hAT2, cluster 1) such as *Acly*, *Hmgcr*, and *Hmgcs1* (Fig. 1D and Fig. S1C). We also found enriched expression of genes induced by an inflammatory response such as *Ptges*, *Lcn1*, *Orm1*, *Tmem173*, and *Ifitm2/3* in this cluster (Fig. 1D and Fig. S1C) (Fortier et al., 2008; Kuriakose and Kanneganti, 2018; Ligresti et al., 2012). Remarkably, essential regulators for AT2 lineage specification such as *Etv5*, *Abca3*, and *Cebpa* were also downregulated, suggesting that this population had lost AT2 identity (Fig. 1D and Fig. S1C) (Martis et al., 2006; Rindler et al., 2017; Zhang et al., 2017), suggesting ‘primed AT2 state (pAT2)’. In addition, cluster 2 cells had a transcriptional signature similar to that of cAT2 cells with the exception of cell cycle-related genes. We also identified an uncharacterized cellular subset of cluster 4 which we named ‘Damage-Associated Transient Progenitors (DATPs)’. DATPs expressed specific markers such as *Cldn4*, *Krt8*, *Ndrg1*, *Sprr1a*, and *AW112010* (Fig. 1D and Fig. S1C). Overall, DATPs shared features of the AT1 lineage transcription signature, but showed much lower expression of canonical AT1 markers including *Pdpn*, *Hopx*, and *Cav-1* (Fig. 1D and Fig. S1C). Analysis of Gene Ontology (GO) terms further revealed that DATPs were characterized by increased expression of genes associated with p53 signaling (e.g. *Trp53*, *Mdm2*, *Ccnd1, Gdf15*), inhibition of proliferation (e.g. *Cdkn1a*, *Cdkn2a*), hypoxia (*Hif1a*, *Ndrg1*), and the interferon-gamma signaling pathway (e.g. *Ifngr1*, *Ly6a*/*Sca-1*, *Irf7*, *Cxcl16*) (Fig. S1D).

As expected, at day 28 post-injury, we observed substantial increases in the mature AT1 and hAT2 populations while cAT2, pAT2, and DATPs were diminished, reflecting return to alveolar homeostasis after injury (Fig. 1C). To better understand the differentiation paths of AT2 cells to AT1 cells during regeneration, we applied partition-based graph abstraction (PAGA, Fig. 1E) and characterized transcriptional programs ordered along pseudotemporal trajectories (Fig. 1F) (Wolf et al., 2019). PAGA shows that AT2 and AT1 cells are connected via a trajectory that includes pAT2 cells and DATPs (Fig. 1E). cAT2 cells were assigned as the population most close to pAT2 cells, suggesting that priming of naïve AT2 cells prior to initiation of differentiation is closely related with a cell cycle event. After excluding cAT2 cells, pseudotime analysis showed that AT2 transitions into AT1 cells via pAT2 cells and DATPs, similar to that what we observed in PAGA (Fig. 1F) (Haghverdi et al., 2016). Taken together, these findings revealed a differentiation trajectory towards AT1 cell fate acquisition that passes through distinct pAT2 and DATP cell states during regeneration.

### IL-1β secreted from interstitial macrophages triggers reprograming of AT2 cells

Given our data showing increased expression of genes associated with the immune response signatures in pAT2 cells, we next asked whether bleomycin injury resulted in inflammation (Fig. S2A). By flow cytometry analysis, we found dynamic changes in macrophage behaviors across injury response and regeneration. At day 7 post injury, the number and frequency of interstitial macrophages (IMs) were significantly increased, whereas the number and frequency of alveolar macrophages (AMs) were decreased (Fig. S2, B-D). These changes were restored to homeostatic levels by day 28, indicating resolution of acute inflammation. Because macrophages localized near AT2 lineage-labeled cells during acute injury (Fig. S2E), we hypothesized that macrophages may affect the behavior of lineage-labeled cells in response to injury. Importantly, we observed that 3D organoid co-cultures in which AT2 cells were cultured together with IMs in the presence of stromal cells revealed more and larger organoid formation than when they were co-cultured with AMs (Fig. 2A-C) (Lee et al., 2014). To further address the contribution of macrophages in alveolar regeneration after injury, we analyzed scRNA-seq of non-lineage-labeled cells from *SPC-Cre^ERT2^;R26R^tdTomato^* mice, including immune cells, isolated in parallel with samples (PBS, D14 and D28) in Fig. 1 (Fig. S2, F-H). The expression level of *IL-1β*, which is specifically detected in macrophages, was increased at day 14 post injury and decreased to homeostatic levels at day 28 (Fig. S2, H and I). Quantitative PCR (qPCR) analysis on isolated AMs and IMs from uninjured lungs revealed that *IL-1β* is highly and specifically expressed in IMs while *IL-18* is enriched in AMs, consistent with previous reports (Fig. S2J) (Misharin et al., 2017). Furthermore, GM-CSF activation specifically augmented *IL-1β* expression in IMs but did not affect *IL-18* expression in AMs (Fig. S2J). Notably, bleomycin injury stimulated *IL-1β* expression in IMs *in viv*o (Fig. S2K). IL-1β treatment was also sufficient to increase the number and size of organoids formed by AT2 cells (Fig. 2, D and E).

**Figure 2.**
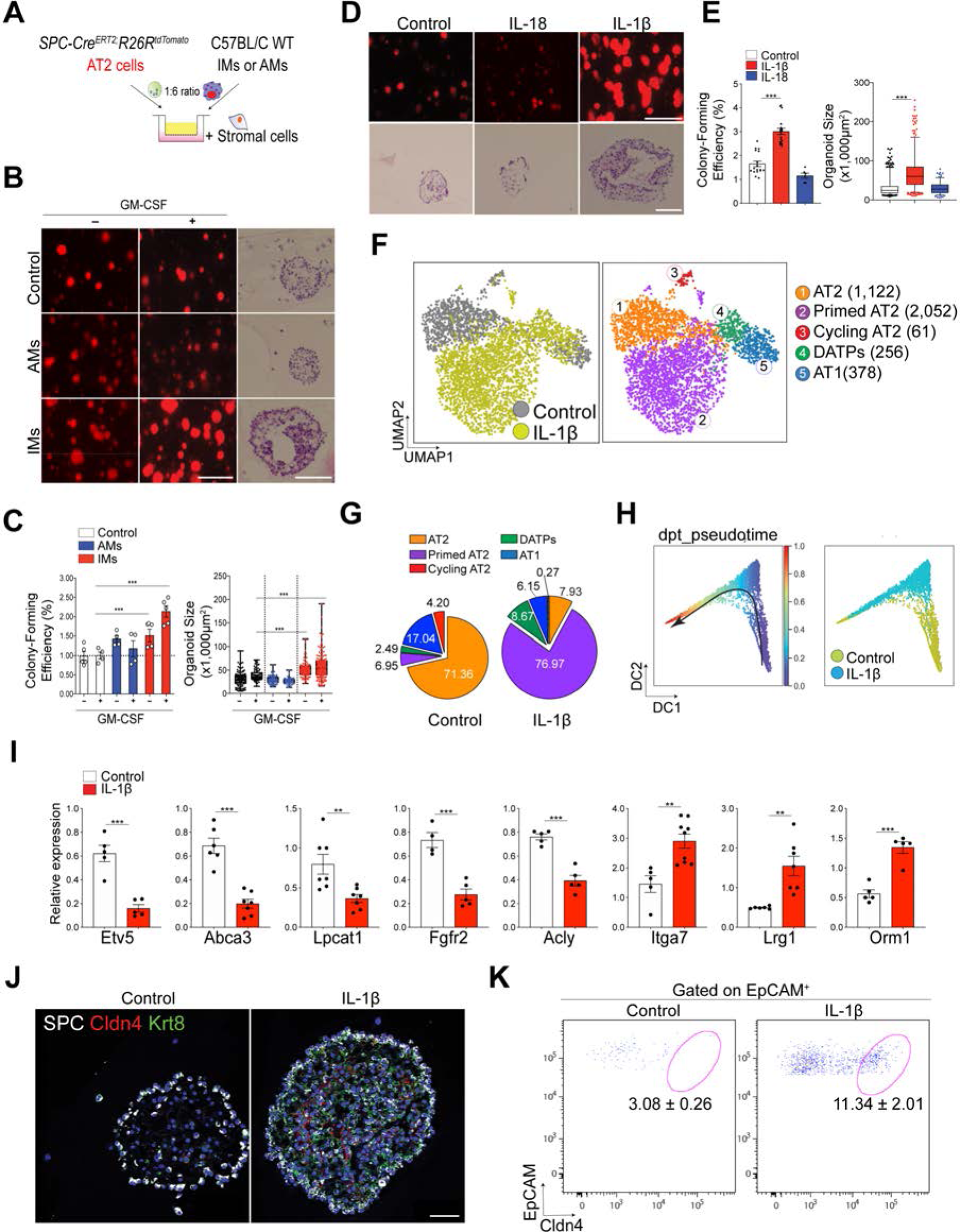
IL-1β signaling directly promotes reprograming of AT2 cells. **(A)** Schematics of organoid co-culture of *SPC* lineage-labeled AT2 cells (SPC^+^Tomato^+^) with interstitial (IMs) or alveolar macrophages (AMs) isolated from wildtype lung tissues in the presence of stromal cells. **See also** Fig. S2. **(B)** Representative fluorescent images (left and middle) and H&E (right) staining of AT2 organoids. GM-CSF was added to activate macrophages. Scale bar, 1,000 μm (left) and 50 μm (right). **(C)** Statistical quantification of colony forming efficiency and size of organoids. Each individual dot represents one experiment from one mouse and date are presented as mean and SEM. ***p<0.001. **(D)** Representative fluorescent images (top) and H&E staining (bottom) of primary organoids derived from *SPC* lineage-labeled AT2 cells (SPC^+^Tomato^+^) treated with vehicle (PBS), IL-1β or IL-18. Scale bar, 1,000 μm (top) and 50 μm (bottom). **(E)** Quantification of colony forming efficiency and size. Date are presented as mean and SEM. **(F)** UMAP visualization of cell clusters from scRNA-seq analysis of epithelial cells from control (1,286 cells) or IL-1β-treated organoids (10 ng/ml, 2,584 cells). Cells were isolated at day 21 in organoid culture. Colors indicate samples and distinct cell types. Number of cells in the individual cluster is depicted in the figure. **See also** Fig. S3. **(G)** The percentage of each cluster in total cells of control or IL-1β-treated organoids. **(H)** Diffusion map according to diffusion pseudotime (DPT, left) order colored by samples (right). **(I)** qPCR analysis of genes that are upregulated (*Itga7*, *Lrg1*, *Orm1*) or downregulated (*Etv5, Abca3, Lpcat1, Fgfr2, and Acly*) in Primed AT2 cells. EpCAM^+^ epithelial cells were isolated from organoids treated with PBS or IL-1β at day 6 in AT2 organoid culture. Each individual dot represents one experiment and data are presented as mean ± SEM. **p<0.01, ***p<0.001. **(J)** Representative IF images showing the generation of DATPs marked by Cldn4 and Krt8 expression in AT2 organoids treated with IL-1β: SPC (white), Cldn4 (red), Krt8 (green) and DAPI (blue). Scale bars, 50 μm. **(K)** Flow cytometry analysis of DATPs by gating with Cldn4 and EpCAM. Data are presented as mean ± SEM (n=5). ***p<0.001.

To further ask how IL-1β affects the cellular and molecular behaviors of AT2 cells, we performed scRNA-seq of control and IL-1β-treated organoids. Based on the marker gene expression, we identified five distinctive clusters (AT2, pAT2, cAT2, DATPs, and AT1 cells) similar to those we had seen in AT2 lineage-labeled cells (Fig. 2F and Fig. S3, A-C). In control organoids, most cell types corresponded to AT2 and AT1 cells alongside smaller pAT2 and DATPs clusters (Fig. 2G). In contrast, IL-1β-treatment increased the pAT2 fraction to ~77% of pAT2 cells, classified by low expression of genes, such as *Etv5*, *Abca3*, and *Cebpa*, suggesting that IL-1β triggers AT2 cells to enter a primed state (Fig. 2G). The DATP population was also increased by IL-1β treatment (Fig. 2G). Pseudotime and PAGA analysis of the scRNA-seq data showed that IL-1β-treated organoids skew differentiation of AT2 cells towards AT1 fate (Fig. 2H and Fig. S3D), by enhancing differentiation into pAT2 and DATP states similar to those of regenerating AT2 cells *in vivo* (Fig. S3, E and F). To investigate if IL-1β directly influences AT2 cell fate transitions, we examined cellular states at day 6 and 14, two key differentiation time points across organoid formation. At day 6, qPCR analysis of IL-1β-treated organoids showed an enriched transcriptional signature of pAT2-state relative to control organoids (Fig. 2I). In addition, day 14 immunostaining and flow cytometry analysis for DATP markers, such as Krt8 and Cldn4, confirmed that DATPs were significantly increased in IL-1β-treated organoids (Fig. 2, J and K). These data show that IL-1β treatment in AT2 organoids recapitulate key aspects of *in vivo* lung regeneration. Taken together, our data demonstrate that an IL-1β-mediated inflammatory niche triggers AT2 mediated injury response during alveolar regeneration via proceeding differentiation programs to generate DATPs.

### DATPs differentiate into AT1 and AT2 cells during alveolar regeneration after injury

Our scRNA-seq analysis revealed the previously unknown AT2-lineage derived DATP population emerging during alveolar regeneration and in organoids stimulated with IL-1β. Using AT2 reporter mice (*SPC-Cre^ERT2^;R26R^tdTomato^*), we found that approximately 10% of AT2 lineage-labeled cells express Krt8 at 14 days after bleomycin injury, confirming that DATPs originate directly from AT2 cells (Fig. 3, A-C). Importantly, neither the AT2 marker SPC, nor the AT1 marker Pdpn were detected in this population (Fig. 3, B and D). To further assess functional contributions of DATPs to alveolar regeneration, we established lineage reporter mice for N-Myc Downstream Regulated 1 (*Ndrg1*) which is uniquely expressed in DATPs during alveolar regeneration (*Ndrg1-Cre^ERT2^;R26R^tdTomato^*) (figs. 1D and 3E). We did not detect any expression of *Ndrg1* in airway epithelial cells with or without injury (Fig. S4, A and B). Consistent with our scRNA-seq data, neither AT2 and AT1 cells were labeled by *Ndrg1* expression in PBS control mice (Fig. 3, I and K, and Fig. S4, C-E). However, at 9 days after bleomycin injury, *Ndrg1* lineage-labeled cells emerged with a majority of cells positive for Krt8 in the alveolar region (Fig. 3, F and G). At day 28, we found that approximately 30% of AT1 cells were lineage-labeled by *Ndrg1* with AT1 cell morphology (Fig. 3, J and K). We also confirmed the contribution of DATPs in AT1 cell generation with lineage-tracing analysis using *Krt8* reporter mice (*Krt8-Cre^ERT2^;R26R^tdTomato^*) (Fig. S5A). Consistent with *Ndrg1* lineage-labeled cells, neither AT2 nor AT1 cells were labeled in uninjured lungs (Fig. S5B). *Krt8* expression was only detected in Cldn4^+^ DATPs at day 9 in the alveolar region post injury, but was then prominent in Pdpn+ AT1 cells at day 28 post injury (Fig. S5, C-F).

**Figure 3.**
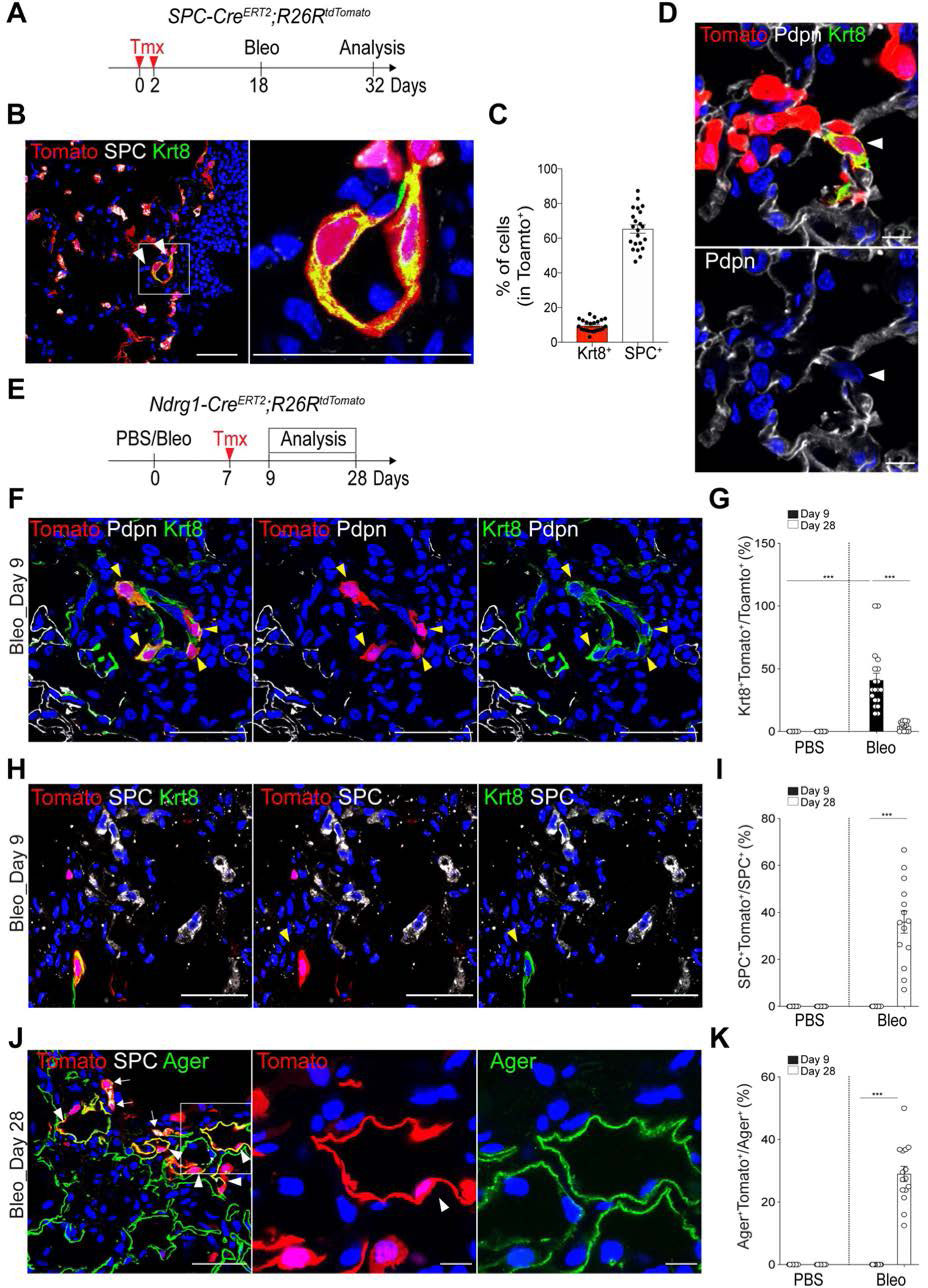
Injury response-specific DATPs are derived from AT2 cells and mediate AT1 lineage differentiation. **(A)** Schematics of experimental design for *SPC* lineage-tracing analysis using *SPC-Cre^ERT2^;R26R^tdTomato^* mice at indicated time points after bleomycin injury. **(B)** Representative IF images showing the derivation of DATPs from AT2 lineage-labeled cells at day 14 post injury: Tomato (red), SPC (white), and Krt8 (green). White boxed insets are shown on the right. Arrowhead points to lineage-labeled *Krt8*^+^DATPs that do not express AT2 marker SPC. Scale bar, 50 μm. **(C)** Quantification of lineage-labeled SPC^+^ AT2 cells or *Krt8*^+^ DATPs at day 14 post injury. Each individual dot represents one section and data are presented as mean ± SEM with three independent experiments (n=4). **(D)** Representative IF images showing the derivation of DATPs from AT2 lineage-labeled cells at day 14 post injury: Tomato (red), Pdpn (white). Arrowhead points to lineage-labeled *Krt8*^+^ DATPs that do not express AT1 marker Pdpn. Scale bar, 10 μm. **(E)** Experimental design for *Ndrg1* lineage-tracing analysis using *Ndrg1-Cre^ERT2^;R26R^tdTomato^* mice after bleomycin injury. Specific time points for tamoxifen injection and analysis are indicated. **See also** Fig. S4. **(F)** Representative IF images showing the derivation of *Ndrg1* lineage-labeled DATPs that are negative for AT1 marker but positive for Krt8 at day 9 post injury: Tomato (red), Pdpn (white), Krt8 (green), and DAPI (blue). Arrowhead points to lineage-labeled DATPs. Scale bar, 50 μm. **(G)** Statistical quantification of Krt8^+^Tomato^+^ cells at indicated time points post PBS or bleomycin injury. Each individual dot represents one section and data are presented as mean ± SEM (n=2 PBS control, n=3 for bleomycin). ***p<0.001. **(H)** Representative IF images showing the derivation of *Ndrg1* lineage-labeled DATPs that are negative for AT2 marker at day 9 post injury: Tomato (red), SPC (white), Krt8 (green), and DAPI (blue). Arrowhead points to lineage-labeled DATPs. Scale bar, 50 μm. **(I)** Statistical quantification of lineage-labeled AT2 (SPC^+^Tomato^+^) cells at indicated time points post PBS or bleomycin injury. Each individual dot represents one section and data are presented as mean ± SEM (n=2 PBS control, n=3 for bleomycin). ***p<0.001. **(J)** Representative IF images showing the differentiation of *Ndrg1* lineage-labeled AT1 and AT2 cells at day 28 after injury: Tomato (red), SPC (white), Ager (green), and DAPI (blue). Arrowhead points to lineage-labeled Ager^+^ cells. White boxed insets (left) are shown on the right. Scale bar, 50 μm (left) and 10 μm (right). **(K)** Statistical quantification of Ager^+^Tomato^+^ AT1 cells at indicated time points post PBS or bleomycin injury. Each individual dot represents one section and data are presented as mean ± SEM (n=2 PBS control, n=3 for bleomycin). ***p<0.001. **See also** Fig. S5.

We observed that a significant number of SPC^+^ AT2 cells were lineage-labeled by *Ndrg1* and *Krt8* at day 28 post bleomycin injury (Fig. 3, I and J and Fig. S5, G and H). To confirm that DATPs possessed capacity of dedifferentiating into AT2 cells, we isolated AT2 cells (CD31^−^CD45^−^EpCAM^+^MHCII^+^) (Hasegawa et al., 2017) from *Krt8* reporter mice and performed organoid cultures in the presence of IL-1β (Fig. S5I). At day 14 in culture, we added 4-OH tamoxifen to label *Krt8*-expressing DATPs. Consistent with immunostaining for Krt8 in organoids (Fig. 2J), we detected Tomato^+^ cells (*Krt8*^+^ DATPs) in the inner part of organoids, which segregated distinctly from Tomato^−^MHCII^+^ AT2 cells by flow cytometric analysis (Fig. S5, J and K). Furthermore, *Krt8*+ DATPs (Tomato^+^MHCII^−^) isolated from organoids were capable of forming organoids composed of DATPs and SPC+ AT2 cells (Fig. S5, K-M).

### IL-1β signaling is required for cell fate conversion into DATPs during alveolar regeneration

Given that IL-1β treatment increased generation of DATPs in organoids, we next asked whether IL-1β signaling is required for differentiation into DATPs *in vivo*. To answer this question, we generated *Il1r1*^flox/flox^*;SPC-Cre^ERT2^;R26R^tdTomato^* mice to deplete *Il1r1*, a functional receptor for IL-1β, specifically in AT2 cells (Fig. 4A). The proliferative activity of *Il1r1*-deficient AT2 cells was comparable to that of *Il1r1*-haplodeficient AT2 cells post injury (Fig. S6A). As IL-1β treatment increased organoid size and forming efficiency (Fig. 2, D and E), we carefully examined AT2 cell proliferation by EdU incorporation assays at early time point (day 4) in organoid cultures. Although IL-1β-treated organoids revealed increases in EdU incorporation rates relative to control, notably, *Il1r1*-deficient AT2 cells also showed a similar rate of EdU incorporation, indicating that IL-1β does not directly influence on AT2 cell proliferation (Fig. S6B). Given that differential expressions of growth factors regulating AT2 cell proliferation in IL-1β-treated stromal cells co-cultured with AT2 cells in organoids (Fig. S6, C-E), it is highly likely that IL-1β enhances AT2 cell proliferation via modulating surrounding cells rather than direct effects on AT2 cells.

**Figure 4.**
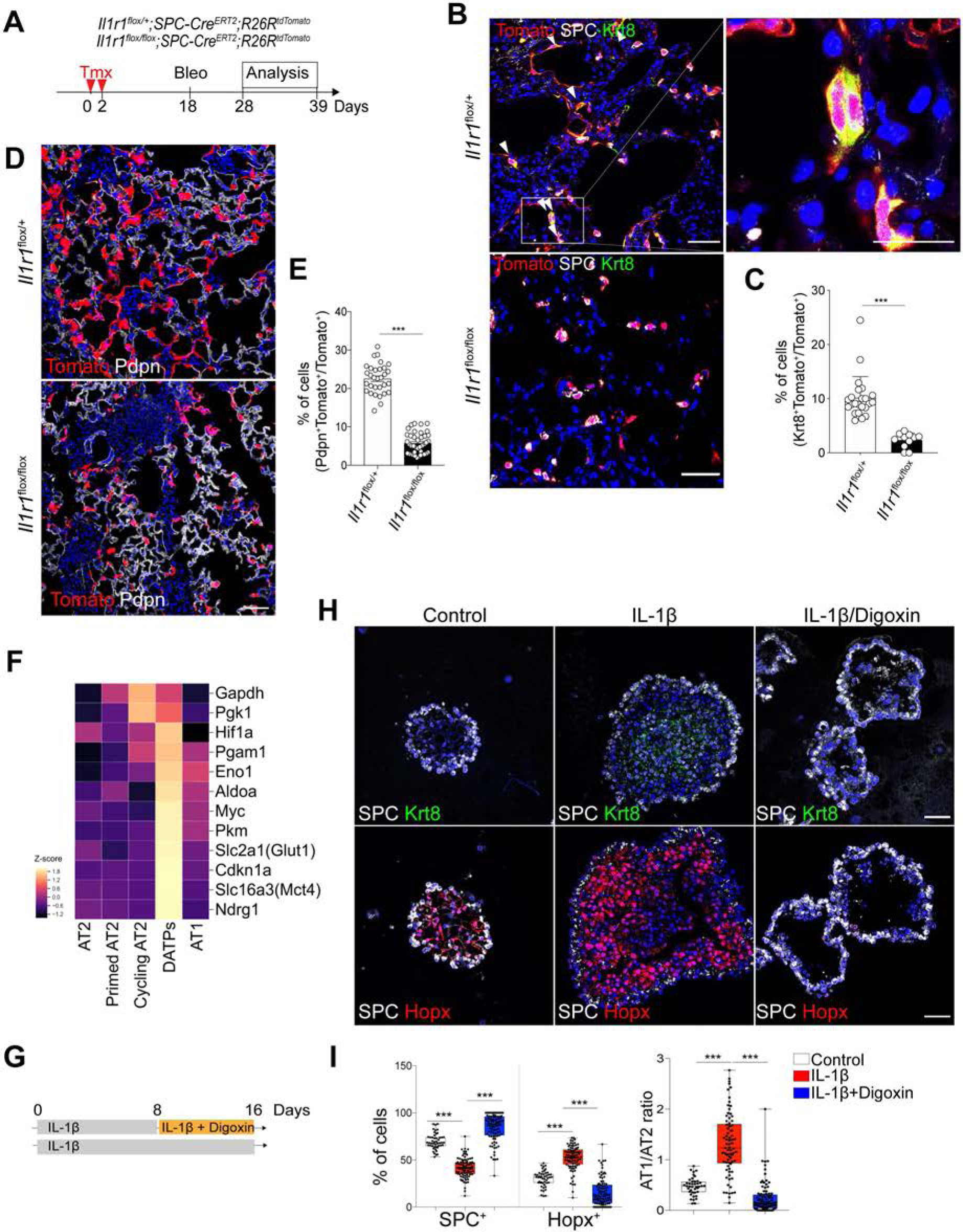
DATPs induced by IL-1β-driven Hif1a signaling are essential mediators for alveolar regeneration. **(A)** Experimental design for lineage tracing of *Il1r1*-haplodeficient or deficient AT2 cells post bleomycin administration. **(B)** Representative IF images showing DATPs generation from *SPC* lineage-labeled cells at day 10 post injury in the indicated genotype: Tomato (for *SPC* lineage, red), SPC (white), Krt8 (green), and DAPI (blue). Scale bars, 50 μm. **See also** Fig. S6. **(C)** Quantification of lineage-labeled *Krt8^+^* DATPs at day 10 post injury. Each individual dot represents one section and data are presented as mean ± SEM (n=3). **(D)** Representative IF images showing AT1 cell differentiation from *SPC* lineage-labeled cells at day 21 post injury in the indicated genotype: Tomato (for *SPC* lineage, red), Pdpn (white), and DAPI (blue). Scale bars, 50 μm. **(E)** Quantification of lineage-labeled Pdpn^+^ AT1 cells at day 21 post injury. Each individual dot represents one section and data are presented as mean ± SEM (n=6). **(F)** Heat map of the transcriptional profiles of genes that are associated with Hif1a-mediated signaling including glycolysis pathway in the subset of clusters. **See also** Fig. S7. **(G)** Schematic of AT2 organoid culture treated with digoxin in the presence of IL-1β. **(H)** Representative IF images showing the impaired generation of DATPs and AT1 lineage in digoxin-treated organoids: SPC (white), Krt8 (top, green), Hopx (bottom, red), and DAPI (blue). Scale bar, 50 μm. **(I)** Quantification of the frequency of AT2 (SPC^+^) or AT1 (Hopx^+^) cells (left) and the ratio of AT1/AT2 (right). Each individual dot represents one experiment and data are presented as mean ± SEM. ***p<0.001. **See also** Fig. S8.

We then further analyzed cAT2 subsets (derived from AT2 lineage-labeled cells post injury, Fig. 1B), which showed step-wise cell cycle transitions based on expression of cell cycle phase-specific genes (Fig. S6F). We discovered that AT2 cells acquired transcriptional signatures of pAT2 cells during the transition from S to G2/M phase in the cell cycle (Fig. S6G). During this transition, the expression of naïve AT2 cell markers including *Abca3* was downregulated while the expression of genes associated with inflammatory response including *Ptges* was induced. Remarkably, *Il1r1* expression was upregulated specifically in G2/M phase (Fig. S6G). Importantly, we found that *Il1r1*-deficient AT2 cells failed to differentiate into DATPs at day 10 post injury (Fig. 4, B and C). Subsequently, lineage-labeled AT1 cells were significantly decreased at day 21 post injury, indicating impaired differentiation of AT2 cells into AT1 cells in the absence of IL-1β signaling (Fig. 4, D and E). Overall, these findings suggest that IL-1β does not directly influence proliferative properties of AT2 cells, but instead primes AT2 cells to initiate cell fate transition into DATPs during alveolar regeneration.

### Hif1a signaling is integral for DATP cell conversion and AT1 differentiation

In our next set of experiments, we asked which downstream targets/factors driven by IL-1β are required for DATP differentiation. Upon further analysis of our *in vivo* and *in vitro* scRNA-seq data, we discovered a unique metabolic signature with higher expression of genes involved in glycolysis pathway such as *Pgk1*, *Pkm*, and *Slc16a3* (Fig. 4F). By measuring the extracellular acidification rate (ECAR) in organoids, we found that IL-1β enhanced the glycolysis metabolism (Fig. S7, A-C). IL-1β-treated organoids also showed higher rates of glucose uptake compared to control (Fig. S7, D and E). Notably, expression of *Hif1a*, a critical regulator for aerobic glycolysis metabolism, was enriched in DATPs (Fig. 4F) (Dang et al., 2008; Semenza, 2012). To determine whether Hif1a signaling is required for the transition into DATPs, we treated AT2 organoids with digoxin, a potent inhibitor of Hif1a activity, in the presence of IL-1β (Fig. 4G). At day 6 in culture, when higher gene signatures of pAT2 cells were detected, digoxin-treated organoids showed impaired generation of DATPs and AT1 cells (Fig. 4, H and I). We next deleted *Hif1a* specifically on AT2 cells using *Hif1a*^flox/flox^*;SPC-Cre^ERT2^;R26R^tdTomato^* mice (Fig. S8A). Consistent with our observations in organoid results, *Hif1a*-deficient AT2 cells failed to generate DATPs at day 10 post injury (Fig. S8, B and C). Similarly to *Il1r1*-deficient AT2 cells, AT2 cells lacking *Hif1a* failed to differentiate into AT1 cells (Fig. S8, D and E). Taken together, these results demonstrate that IL-1β enhances Hif1a-mediated glycolysis metabolic changes which are integral for the transition into DATPs and subsequent differentiation into AT1 cells during injury repair.

### *Il1r1*^+^AT2 cells are functionally and epigenetically distinct subsets that generate DATPs by IL-1β signals in alveolar regeneration

Given the importance of IL-1β signaling in alveolar regeneration, we asked whether all AT2 cells are equally capable of responding to IL-1β inflammatory signals. To answer this question, we generated *Il1r1* reporter mice (*Il1r1-Cre^ERT2^;R26R^tdTomato^*) and treated them with tamoxifen to lineage trace *Il1r1*-expressing cells (Fig. 5, A and B). We found that *Il1r1* was expressed in airway ciliated cells and small subsets of mesenchyme cells in uninjured lungs (Fig. 5C and Fig. S9). Although ~ 15% of AT2 cells were lineage-labeled in the uninjured lung, bleomycin injury significantly increased the population of lineage-labeled AT2 cells up to ~60% at day 14 post injury (Fig. 5, C and D). *Il1r1* lineage-labeled AT2 cells were also more proliferative than unlabeled AT2 cells (Fig. 5, E and F). Approximately 80% of DATPs were lineage-labeled by *Il1r1*, suggesting that DATPs are mainly originating from *Il1r1+*AT2 cells (Fig. 5, G and H). At day 28 post injury, lineage-labeled AT1 cells were nicely observed (Fig. 5I).

**Figure 5.**
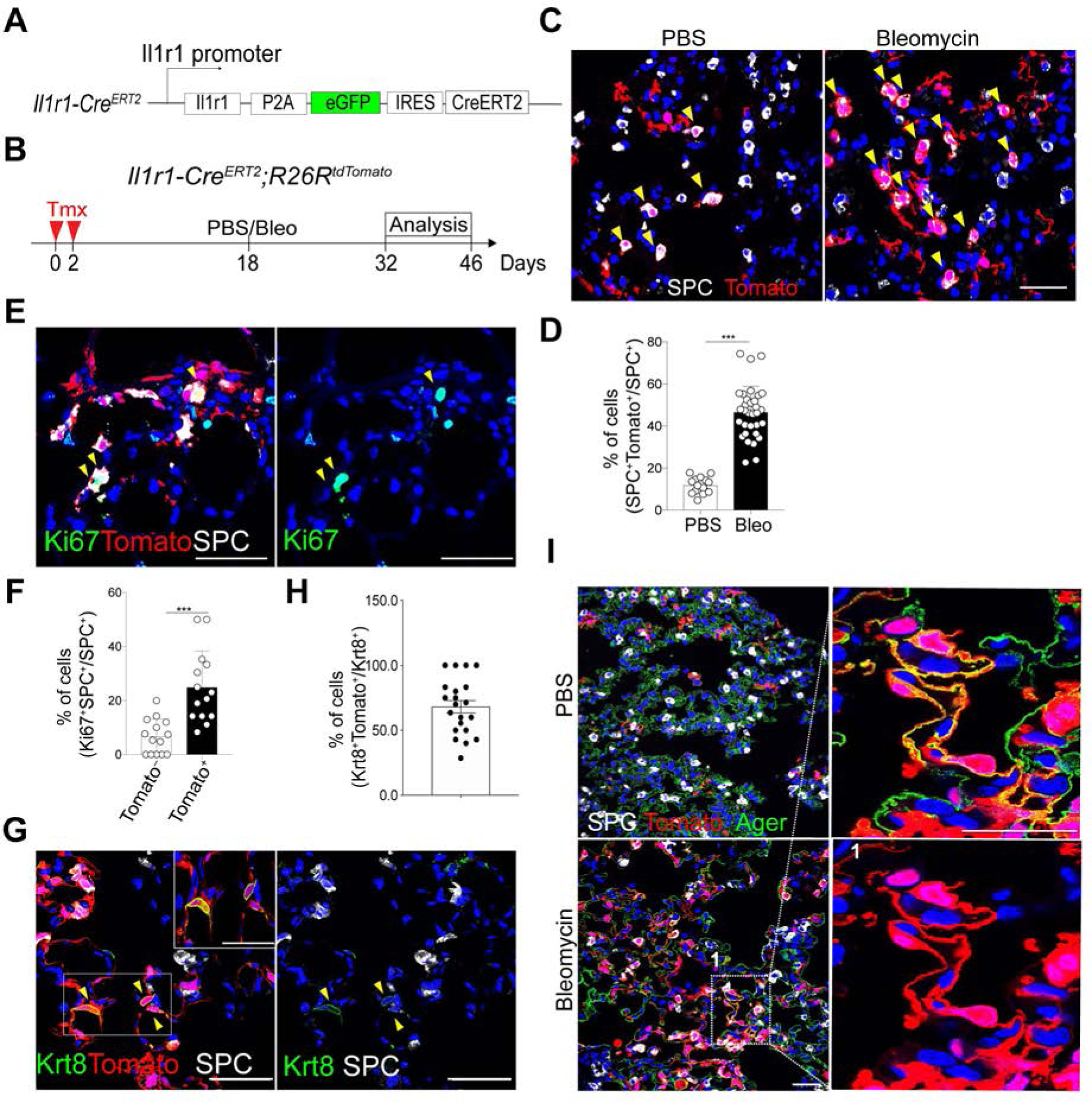
*Il1r1*^+^AT2 cells are distinct subsets that generate DATPs during alveolar regeneration after injury. **(A)** Schematic of *Il1r1-CreERT2* mice. See also Fig. S9. **(B)** Experimental design for lineage tracing. Date for analysis is as indicated. **(C)** Representative IF images showing *Il1r1* lineage-labeled AT2 cells in the lung of mice treated with control (PBS) or bleomycin at day 14 post injury: Tomato (for *Il1r1* lineage, red), SPC (white), and DAPI (blue). Arrowheads, *Il1r1* lineage-labeled SPC^+^AT2 cells. Scale bars, 50 μm. **(D)** Quantification of *Il1r1* lineage-labeled SPC^+^ AT2 cells in (C). Each individual dot represents one section and data are presented as mean ± SEM with three independent experiments. ***p<0.001. **(E)** Representative IF images showing Ki67^+^ cells in lineage-labeled or–unlabeled SPC^+^ AT2 cells at day 14 post injury: Tomato (for *Il1r1* lineage, red), SPC (white), Ki67 (green), and DAPI (blue). Arrowheads, *Il1r1* lineage-labeled proliferating AT2 cells. Scale bars, 50 μm. **(F)** Quantification of Ki67^+^ AT2 cells in lineage-labeled or-unlabeled SPC^+^ cells. Each individual dot represents one section and data are presented as mean ± SEM with three independent experiments. ***p<0.001. **(G)** Representative IF images showing *Il1r1* lineage-labeled DATPs at day 28 post injury: Tomato (for *Il1r1* lineage, red), Krt8 (green), and DAPI (blue). Arrowheads, *Il1r1* lineage-labeled DATPs. Insets (left) show high-power view (right top). Scale bars, 50μm. **(H)** Quantification of *Il1r1* lineage-labeled DATPs at day 14 post bleomycin injury. Each individual dot represents one section and data are presented as mean ± SEM with three independent experiments. **(I)** Representative IF images showing *Il1r1* lineage-labeled AT1 cells at day 28 post injury: Tomato (for *Il1r1* lineage, red), SPC (white), Ager (green), and DAPI (blue). Scale bars, 50μm.

We posited that epigenetic mechanisms might shape the active response of *Il1r1^+^*AT2 cells and next performed ATAC-seq (Assay for Transposase-Accessible Chromatin with high-throughput sequencing). Although most gene including AT2 markers and general housekeeping genes showed similar chromatin accessibility patterns, notable differences were present in the open chromatin states in *Il1r1^+^*AT2 cells relative to bulk AT2 cells (Fig. 6, A and B and Fig. S10). Analysis for Gene Ontology (GO) terms distribution of highlighted genes revealed that epigenetic regulation and inflammation-associated pathways including Interleukin-1 signaling were enriched in *Il1r1^+^*AT2 cells (Fig. 6, C and D). Motif analysis of DNA binding-site showed that *Il1r1^+^*AT2-enriched chromatin contains motifs for key transcriptional factors associated with inflammation such as AP-1, CREB, NF-kB and Rorc, while shared genes were enriched in motifs for key lung development factors as Nkx2.1 and Cebp (Fig. 6E) (Martis et al., 2006; Minoo et al., 1999; Miossec and Kolls, 2012; Schonthaler et al., 2011). Taken together, these results demonstrate that *Il1r1* marks epigenetically distinct AT2 cell subtypes with capacity for rapid expansion and subsequent differentiation into AT1 cells during injury response.

**Figure 6.**
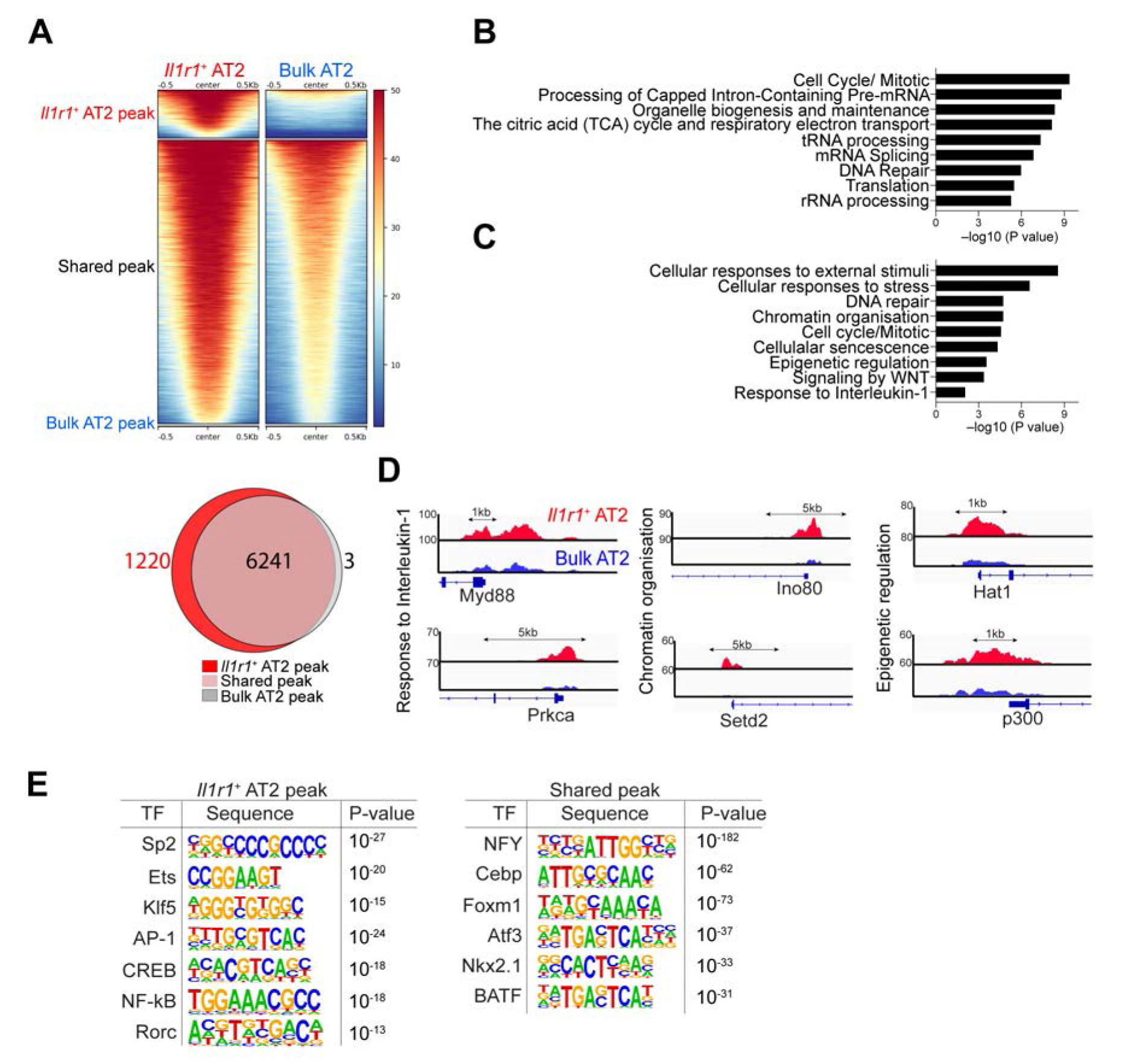
*Il1r1*^+^AT2 cells possess a chromatin architecture that enables a rapid response to injury. **(A)** ATAC-seq heat map (Top) and Venn diagrams (bottom) showing genome-wide regions of differential open chromatin peaks in *Il1r1*^+^AT2 versus bulk AT2 cells in duplicates. The values correspond to the peak signal distribution around TSS (Transcription Start Sites). Number of nearest neighbour genes covered by peaks are indicated on diagrams. **(B)** GO enrichment analysis of the nearest neighbour genes in the vicinity of peaks shared between *Il1r1*^+^AT2 and bulk AT2 cells. **(C)** GO enrichment analysis of the nearest neighbour genes in the vicinity of *Il1r1*^+^AT2 peaks. **(D)** Snapshots of genomic loci in which the chromatin-accessible peaks are specifically opened in *Il1r1*^+^AT2 cells identified by GO enrichment analysis shown in (**C**). **(E)** Transcription factor motif enrichment within *Il1r1*^+^AT2-specific peaks or peaks shared between *Il1r1*^+^AT2 and bulk AT2 cells. see also Fig. S10.

### Chronic inflammation mediated by sustained IL-1β levels stalls transition of DATPs into mature AT1 cells

Although expression levels of early AT1 markers such as *Lmo7*, *Pdpn*, and *Hopx* were comparable in control and IL-1β-treated organoids (Fig. S11A), we found that AT1-like cells present in IL-1β-treated organoids failed to upregulate mature AT1 markers highly expressed in control AT1 cells such as *Aqp5*, *Vegfa*, *Cav-1*, and *Spock2* (Fig. S11B). Instead, AT1-like populations in IL-1β-treated organoids highly expressed DATP-associated genes including *Cldn4*, *AW112010*, and *Lhfp* (Fig. S11C), indicating that sustained IL-1β treatment in AT2 organoids causes accumulation of DATPs and prevents terminally differentiation into mature AT1 cells. We then asked whether the stalled transition to mature AT1 cells could be rescued by relieving IL-1β-mediated inflammation. We cultured AT2 organoids with IL-1β for 14 days and maintained them for an additional 7 days without IL-1β treatment (Fig. 7A). Indeed, we found that expression of late AT1 markers became significantly upregulated upon IL-1β withdrawal (Fig. 7B), concomitant with downregulation of DATP markers and expression of *Hif1a* and other glycolysis pathway genes (Fig. S11D). These findings prompted us ask whether inhibition of glycolysis in stalled DATPs might facilitate AT1 cell maturation. To this end, we treated AT2 organoids with IL-1β for 14 days and then with the glycolysis inhibitor 2-deoxyglucose (2-DG, a glucose analogue that causes hexokinase inhibition and disruption of glycolysis) in the continued presence of IL-1β for additional 4 days (Fig. 7C). Notably, inhibition of high glucose metabolism significantly upregulated expression of mature AT1 makers (Fig. 7D). With immunostaining, we confirmed that AT2 cells with persistent IL-1β treatment failed to generate mature AT1 cells expressing Cav-1, a late AT1 cell marker, whereas the expression level of Hopx, an early AT1 cell marker, was comparable to that seen in control (Fig. 7E). Importantly, 2-DG-treated organoids rescued the impaired maturation of AT1 cells even in the presence of IL-1β (Fig. 7E).

**Figure 7.**
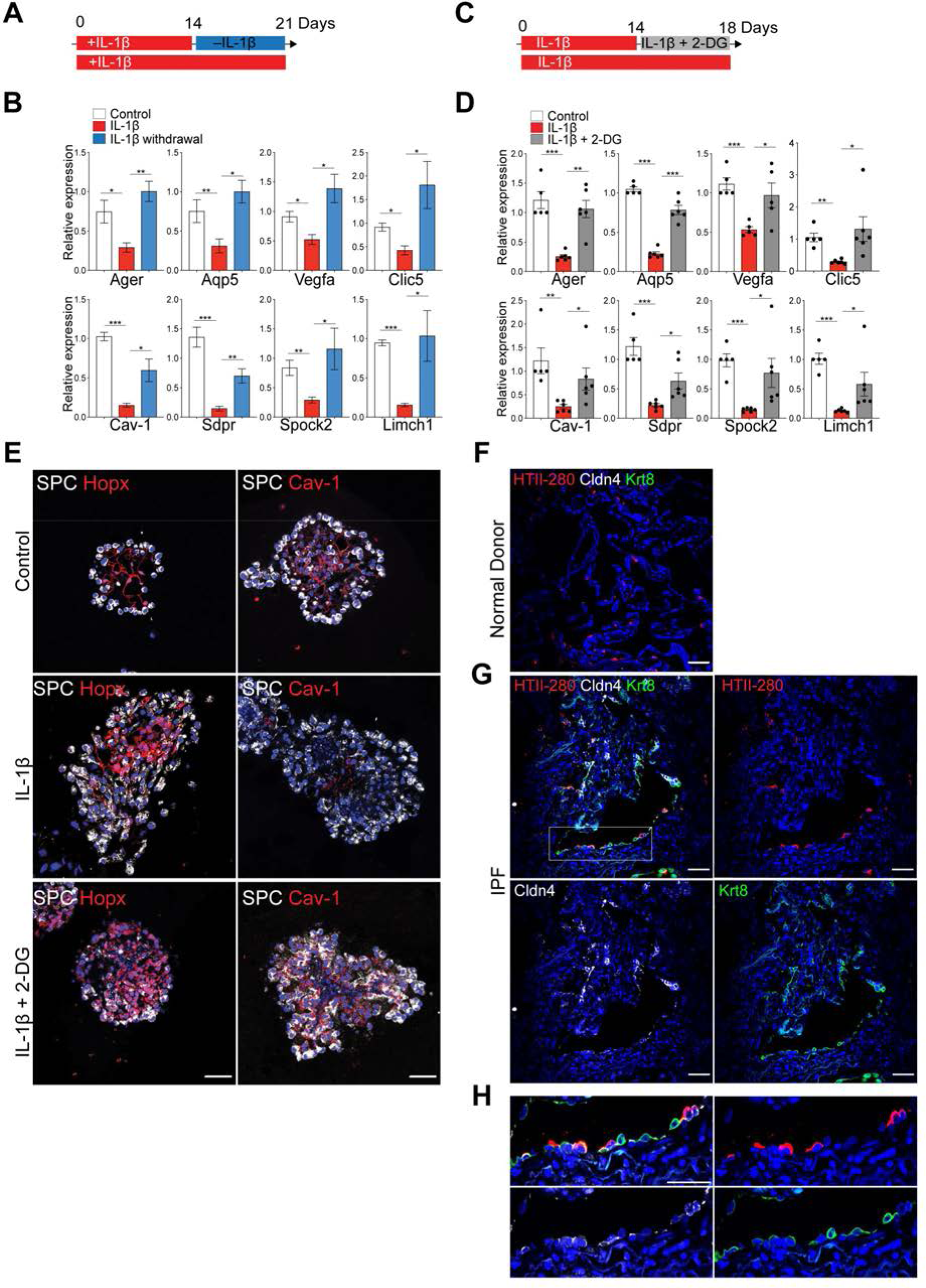
Glycolysis pathway driven by IL-1β prevent DATPs from converting into terminally mature AT1 cells. **(A)** Schematic of AT2 organoid culture treated with or without IL-1β. **(B)** qPCR analysis for mature AT1 markers on isolated epithelial cells from AT2 organoids. Date are presented as mean ± SEM of four biological replicates from two-independent experiments. *p<0.05, **p<0.01, ***p<0.001. **(C)** Schematic of AT2 organoid culture treated with or without 2-deoxy glucose (2-DG) in the presence of IL-1β. **(D)** qPCR analysis for mature AT1 markers on isolated epithelial cells from AT2 organoids. Each individual dot represents one experiment and date are presented as mean ± SEM. *p<0.05, **p<0.01, ***p<0.001. **(E)** Representative IF images showing the rescued maturation of AT1 cells in 2-DG treated organoids in the presence of IL-1β: SPC (white), Hopx (top, red), Cav-1 (bottom, red) and DAPI (blue). Scale bar, 50 μm. See also Fig. S11. **(F)** Representative IF images of KRT8+CLDN4+ DATPs-like population in the lung from normal donors (n=3). HTII-280 (red), CLDN4 (white), KRT8 (green) and DAPI (blue). Scale bar, 50 μm. See also Fig. S12. **(G)** Representative IF images of KRT8+CLDN4+ DATPs-like population in the lung from IPF patients (n=5). HTII-280 (red), CLDN4 (white), KRT8 (green) and DAPI (blue). Scale bar, 50 μm. **(H)** High-power view of white boxed insets in Fig. 7G. HTII-280 (red), CLDN4 (white), KRT8 (green) and DAPI (blue). Scale bar, 50 μm.

We hypothesized that a chronic inflammatory environment will lead to a gradual accumulation of DATPs and eventually defective differentiation and declined lung regeneration. Consistent with our data, recent studies using a high-resolution scRNA-seq analysis reported that a transcriptionally distinct population aberrantly accumulates in a non-permissive pathologic environment such as idiopathic pulmonary fibrosis (IPF) (Adams et al., 2019; Habermann et al., 2019; Kobayashi et al., 2019; Wu et al., 2020). Notably, we also observed abundant KRT8^+^CLDN4^+^ DATPs-like cells next to HTII-280^+^ AT2 cells in alveolar regions of IPF patient tissue samples, but not within the alveoli of normal donor lung (Fig. 7, F-H). In addition, given the close relationship between chronic inflammation and lung cancer, and recent reports suggesting transcriptional features of injury responses in lung tumor cells, we also found that KRT8^+^CLDN4^+^ DATPs-like cells are observed within the tumor in patient tissue samples of lung adenocarcinoma (Fig. S12) (Conway et al., 2016; Mantovani et al., 2008; Maynard et al., 2019; Moll et al., 2018). Taken together, these findings demonstrate that chronic inflammatory signals cause dysregulation of DATPs, which leads to development and/or progression of human lung diseases.

## Discussion

Effectively coordinated tissue repair is critical for maintenance of tissue integrity and function. In responding to environmental assault, the ability to sense physiological changes is essential for stem cells to initiate repair and resolve damage. Here, we focused on how inflammatory stimuli direct the cell fate behavior of AT2 stem cells during lung injury repair. Our data reveals, for the first time, the detailed step-wise differentiation trajectories of AT2 cells, which are regulated by IL-1β-mediated inflammatory signals during the regeneration process. Significantly, we identified *Il1r1*^+^AT2 cells and Damage-Associated Transient Progenitors (DATPs) as two classes of regenerative cell populations dedicated to lung injury repair. Our findings bring new insight into how unresolved inflammation mediated by persistent IL-1β signals prevents cell fate transitions, resulting in impaired regeneration and eventually leading to lung diseases (Fig. 8).

**Figure 8.**
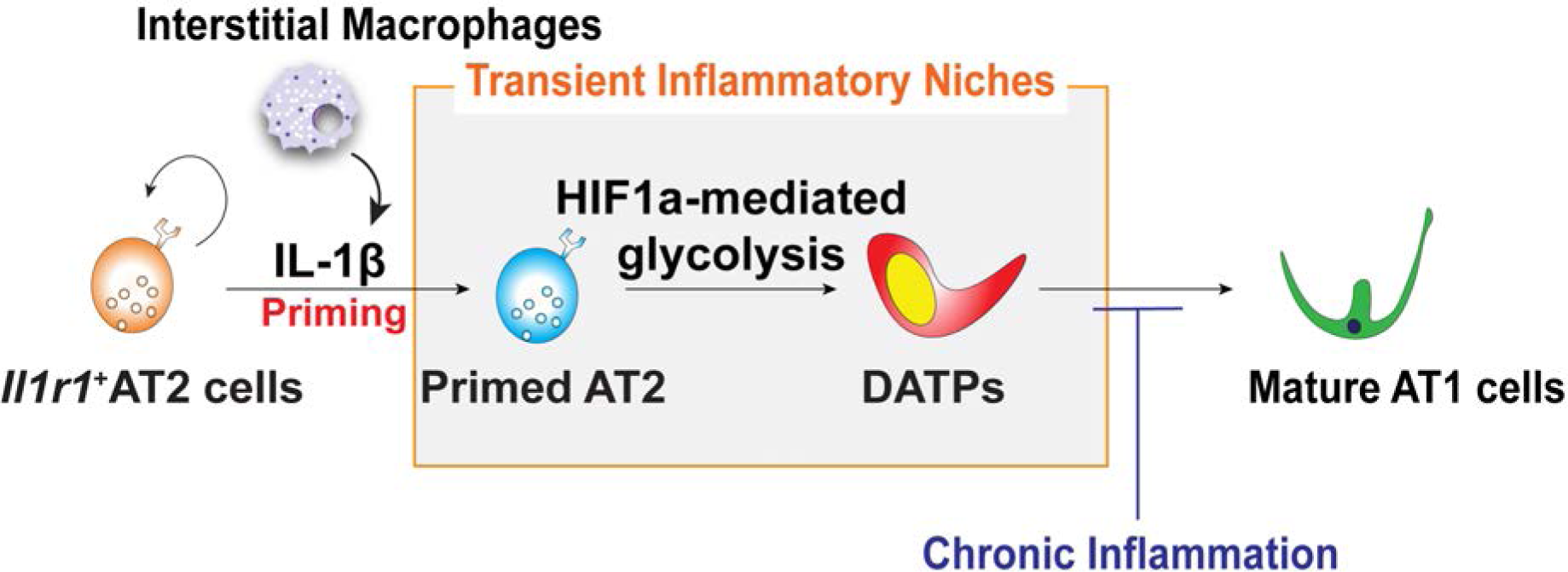
Reconstitution of lineage differentiation programs from AT2 to AT1 cells via DATPs is directed by IL-1β signaling during injury repair. Molecular mechanism governing the AT2 cell reprograming by IL-1β-driven transient inflammatory niches during alveolar regeneration. Pro-inflammatory cytokine IL-1β secreted from interstitial macrophages triggers the reprograming of *Il1r1*^+^AT2 cells to lose AT2 identity and enter a primed state during cell cycle transition. In turn, IL-1β-driven HIF1a signaling promotes the transition into DATPs that are essential mediators for mature AT1 cell differentiation. Moreover, unresolved inflammation mediated by persistent IL-1β signals causes the blockade in the transition to terminally differentiated AT1 lineage, which results in the accumulation of DATPs. This finding highlights the important therapeutic implications of DATPs and their modulation by IL-1β signalling in the context of chronic inflammation associated diseases.

Although mechanisms underlying alveolar regeneration are complex, our scRNA-seq analysis of *in vivo* AT2 lineage-labeled cells and AT2 cell-derived organoids defines the precise reprograming of AT2 cells into AT1 cells during injury repair. We discovered two distinct populations, pAT2 cells and DATPs, as intermediaries between quiescent AT2 and terminally differentiated AT1 lineages. pAT2 cells highly express genes that respond to inflammation (e.g. *Pteges*, *Orm1*, *Zbp1*), are involved in promoting angiogenesis (e.g. *Lrg1*, *Cxcl17*, and *Egfl6/7*) and reduce reactive oxidative species (ROS) (e.g. *Glrx*, *Prdx4*, and *Gstk1/2*). These properties suggest that pAT2 cells actively respond to inflammatory stimuli, reshaping reciprocal interactions between epithelial cells and their niches during tissue repair. pAT2 cells display much lower expression of genes that are essential for AT2 identity and maintenance such as *Etv5* and *Abca3*, while still expressing comparable levels of canonical AT2 markers such as *Sftpc* and *Lyz2*. These transcriptional signatures were also seen in IL-1β-treated AT2 cells, leading us to classify pAT2 cells as a population that is skewed towards the AT1 cell fate.

Our data reveal that pAT2 cells share a transcriptional program resembling that of cAT2 cells but with lower expression levels of cell cycle genes (e.g. *MKi67*, *Cdk1*). Interestingly, we found that transcriptional signatures of pAT2 cells were upregulated during the transition from S to G2/M phase in the cell cycle, suggesting the possibility of entering primed states after exiting proliferation states although further validation studies such as genetic tracing of cAT2 or pAT2 cells are needed to provide the delineated sequence of trajectory between these two states. In addition, at variance with a previous study in *Il1r1*^−/−^ mice (Katsura et al., 2019), we found that proliferative activity of AT2 cells is not directly altered by *Il1r1* depletion in AT2 cells. Instead, our findings in organoid co-culture experiments revealed that stromal cells responding to IL-1β likely support AT2 cell proliferation. scRNA-seq analysis of stromal cells co-cultured with AT2 cells showed that expression of growth factors facilitating AT2 cell proliferation, such as EGFR ligands (e.g. *Ereg*), *Spp1*, and *Hgf* (Ganguly et al., 2014; Zeng et al., 2016) was dramatically increased in IL-1β-treated stromal cells, whereas *Bmp4* (Weaver et al., 2000), which is known to inhibit AT2 cell proliferation was significantly reduced (Fig. S6, C-E). The negative regulators for Bmp4 signaling such as *Grem1/2* were increased in IL-1β-treated stromal cells. Notably, cAT2 cells acquire transcriptional characteristics of pAT2 cells coupled with upregulation of *Il1r1* expression at the transition from S to G2/M phase. These data suggest that IL-1β directly reprograms daughter AT2 cells to enter primed states during the G2/M phase to initiate cell fate transitions without direct influences on cell proliferation. How IL-1β signaling triggers priming of AT2 cells to initiate the differentiation progress remains unknown. Recently, Wnt signaling was reported to prevent reprograming of AT2 cells into AT1 cells (Nabhan et al., 2018), suggesting that crosstalk between IL-1β and Wnt signaling underlies control of cell fate transitions from naïve AT2 to primed cell states.

We discovered a previously unidentified DATP population as an intermediate plastic subpopulation between pAT2 and AT1 cell differentiation states. DATPs expressing *Ndrg1, Cldn4* and *Krt8* are extremely rare at steady-state, yet are significantly induced after injury by IL-1β-mediated inflammatory signaling. Lineage-tracing analysis demonstrated their capacity to give rise to new AT1 cells during alveolar regeneration after injury. Specifically, we determined that IL-1β-driven inflammation and regulation of the Hif1a signaling pathway is essential for DATPs generation. Specific deletion of *Hif1a* in AT2 cells impaired this progression, resulting in deficient production of new AT1 cells. In addition, we also defined that reduction of IL-1β-driven glycolysis is required for transition of DATPs towards initiating AT1 lineage differentiation. This finding suggests that IL-1β-mediated inflammation and transient glycolytic metabolism by generating DATPs may establish a checkpoint determining entry into mature AT1 cell differentiation programs. Of note, DATPs reveal quiescent characteristics represented by expression of cell cycle inhibition, p53 signaling, and senescence marker genes. In addition, emerging evidence supported by a high-resolution scRNA-seq technology suggests an essential role of ‘intermediates’ during developmental process in governing cell fate choices (Olsson et al., 2016). Interestingly, we also found that DATPs may have the plasticity required to revert to the AT2 lineage, in addition to proceeding towards AT1 differentiation.

By combining lineage-trancing and ATAC-seq analysis we uncovered that *Il1r1*^+^AT2 cells take on distinct epigenetic state as they efficiently replenish damaged alveolar lineages in response to IL-1β inflammatory signals. Specific open-chromatin states in regions recognized by epigenetic regulators, including chromatin remodellers (e.g. Ino80) and epigenetic modifiers (e.g. Hat1), allow for their rapid and organized responsiveness to injury during the regeneration process. Significantly, we found that DATPs are mainly arising from *Il1r1*^+^AT2 cells in response to IL-1β signaling after injury. Recently, *Axin2*^+^AT2 cells have been identified as a distinct subset of AT2 cells (Nabhan et al., 2018; Zacharias et al., 2018). Related with the potential role of interconnectivity between IL-1β and Wnt signaling in fate decision of AT2 cells, comparison between *Il1r1*^+^AT2 and *Axin2*^+^AT2 cells will be helpful to understand their relationships during alveolar regeneration.

Resolution of inflammation is a coordinated and active process aimed at restoration of tissue integrity and function. Our data highlight the importance of macrophage activation in the transient inflammatory niche after tissue injury. The increased number of IMs and level of IL-1β peaked at day 14 and resolved to the homeostatic level at day 28 after injury. Analysis of lineage-tracing and scRNA-seq data also revealed that pAT2 cells and DATPs appearing after injury become dramatically reduced as tissue returns to homeostasis. However, significantly, we found that sustained IL-1β signaling causes the defects in the transition from DATPs to terminal differentiation to AT1 lineage, which results in the impaired regeneration. Our finding, for the first time, reveals the cellular and molecular mechanisms how chronic inflammation is implicated in the tissue dysfunction and pathogenesis. Two recent studies showed fibrosis-specific KRT17^+^ cell populations in patient tissues of idiopathic pulmonary fibrosis (IPF) (Adams et al., 2019; Habermann et al., 2019). Here, we find that these populations and DATPs have similar transcriptional signatures, also supported by a recent preprint showing the enriched signatures of *Cldn4^+^* pre-AT1 transitional state in these KRT17^+^ populations in IPF tissues (Kobayashi et al., 2019). Furthermore, we detected KRT8^+^CLDN4^+^ DATPs-like cells in the alveolar regions of IPF tissue samples. In addition, several studies have revealed that mechanisms underlying cancer development co-opt regeneration programs to drive tumoral cellular heterogeneity (Maynard et al., 2019; Moll et al., 2018). Congruent with this work, we also observed DATP-like cells in tissue samples of human lung adenocarcinoma. Our results strongly suggest that fine modulation of DATPs by IL-1β-mediated transient inflammatory niche during injury repair is critical for effective lung restoration and is a potential therapeutic adjunct for treating lung diseases.

### Limitations of the Study

Our study identified subsets of *Il1r1*^+^AT2 cells having distinctive epigenetic signatures and quickly responding to injury-induced inflammation for efficient AT1 cell generation. Despite it is clear that only a subset of AT2 cells expressed *Il1r1* and expanded up to 60% of total AT2 cells during injury repair, we cannot completely rule out the possibility of stochastic expression of cre recombination for *Il1r1* expression during repair process due to the remained tamoxifen activity. Longer wash out periods than 16 days may provide clearer evidences to further define the functionally distinctive subsets of *Il1r1*^+^AT2 cells during injury repair.

## Acknowledgement

We would like to thank Emma Rawlins (University of Cambridge, UK) for valuable scientific discussions and sharing the mouse lines; We would like to thank Randall Johnson (University of Cambridge, UK) for sharing Hif1a^flox/flox^ mouse line; Nisha Narayan and Brian Huntly for sharing materials and discussion on glycolysis experiments; Irina Pshenichnaya (Histology), Maike Paramor (NGS library), Peter Humphreys (Imaging), Andy Riddell (Flow cytometry), Simon McCallum (Flow cytometry, Cambridge NIHR BRC Cell Phenotyping Hub), Katarzyna Kania (single cell sequencing at Cancer Research UK), and Cambridge Stem Cell Institute core facilities for technical assistance; Papworth Hospital Research Tissue Bank for providing tissue samples from IPF and lung adenocarcinoma (T02233); Kelly Evans for sharing histology samples of human lung tissue samples; Seungmin Han and Woochang Hwang for discussion on the scRNA-seq analysis; Life Science Editors for editorial assistance; All Lee Lab members for helpful discussion. This work was supported by Wellcome and the Royal Society (107633/Z/15/Z) and European Research Council Starting Grant (679411). J.C. was supported by the National Research Foundation of Korea (NRF) funded by the Ministry of Education (2017R1A6A3A03005399).

## Author contribution

J.C. and J.-H.L. designed the experiments, interpreted the data, and wrote the manuscript; J.C. performed most experiments and data analysis; J.-E.P. performed and analyzed scRNA-seq data; G.T. and N.H. analyzed ATAC-seq data; M.Y. shared *Ndrg1-* Cre^ERT2^ mouse line; B.-K.K. helped the generation of *Il1r1-*Cre^ERT2^ mouse line.

### STAR METHODS

#### KEY RESOURCES TABLE

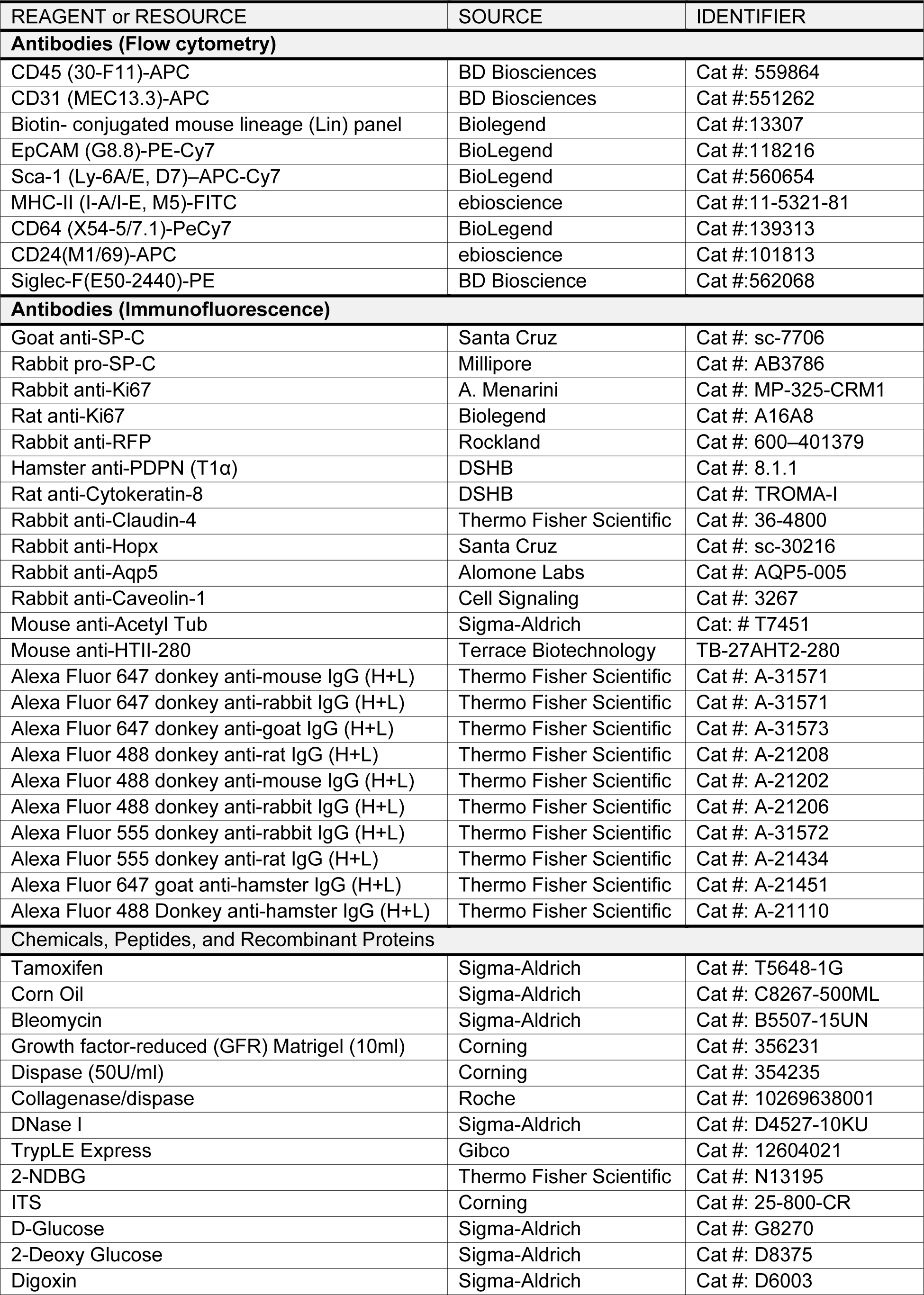
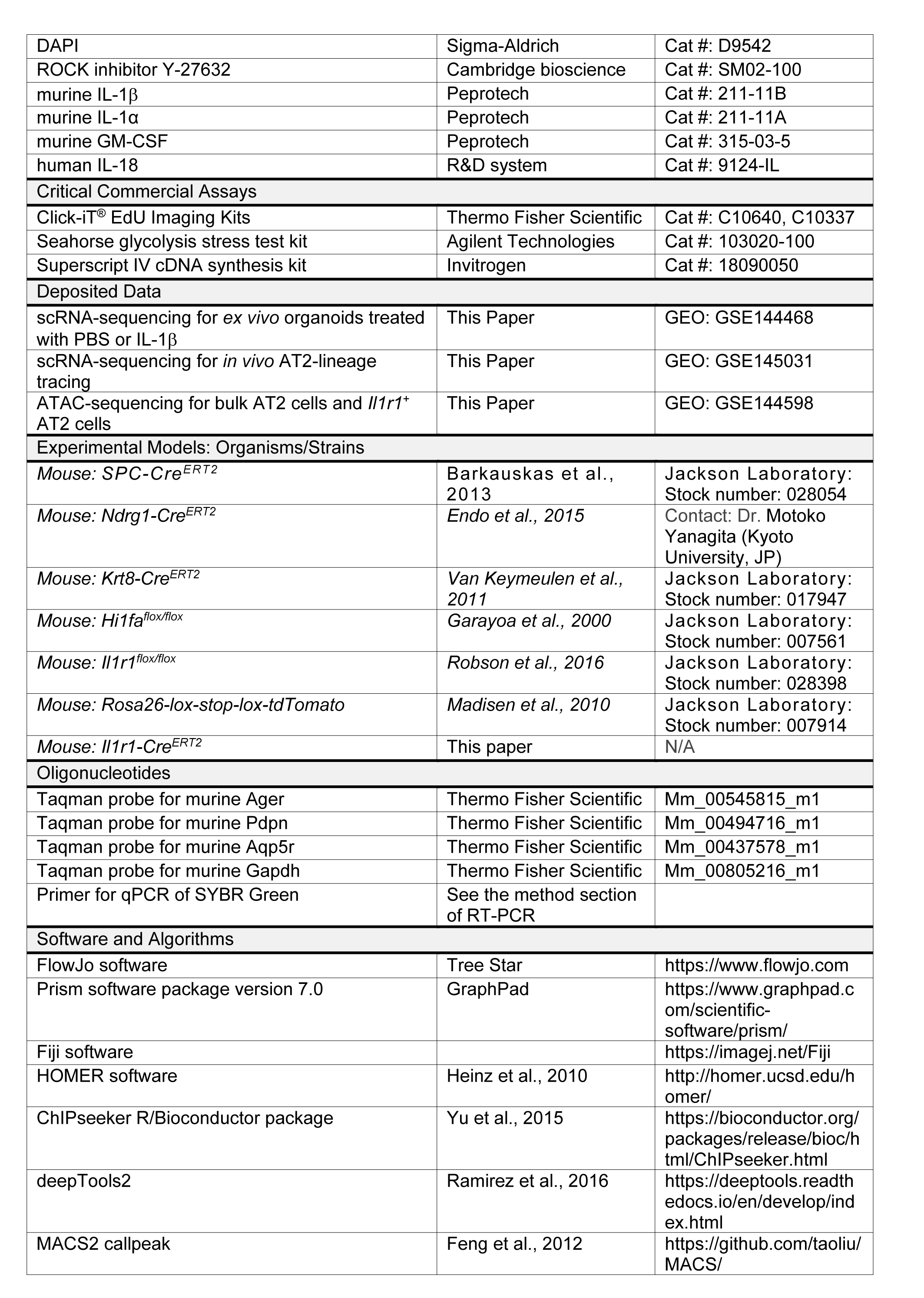
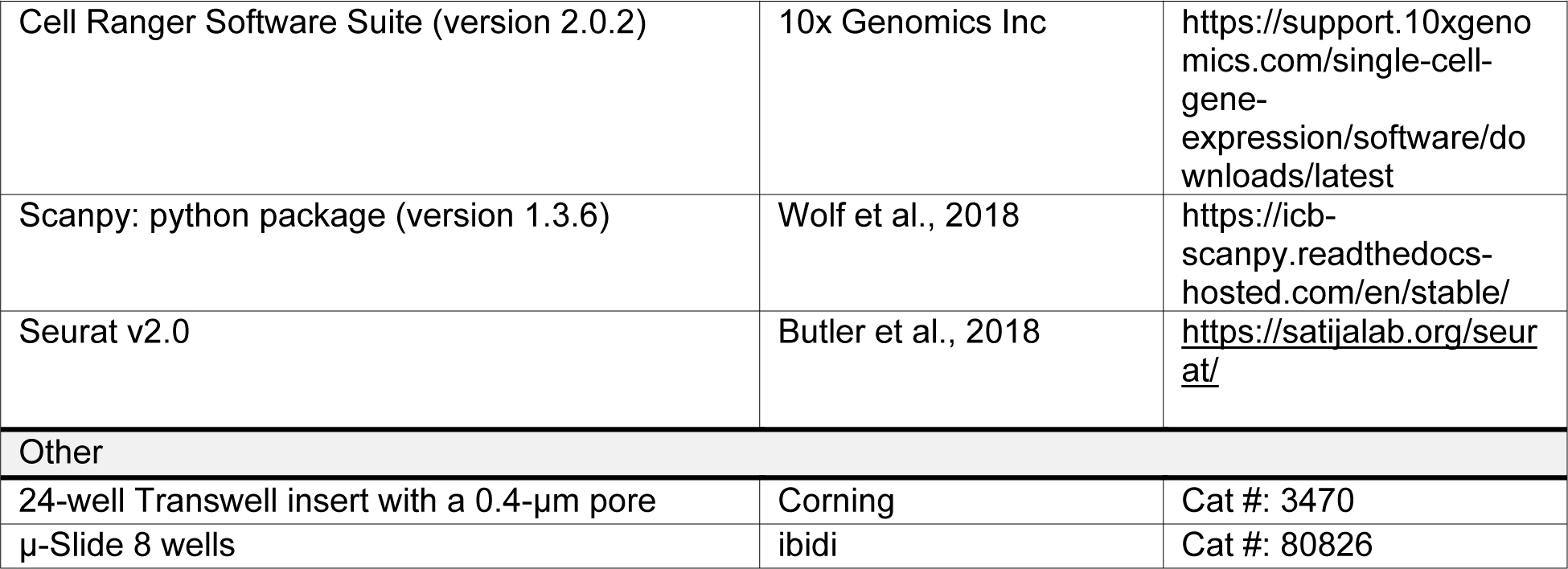

## Methods

### Animals

*SPC-Cre^ERT 2^* (Barkauskas et al., 2013)*, Rosa26-lox-stop-lox-tdTomato* (Madisen et al., 2010), *Ndrg1-Cre^ERT2^* (Endo et al., 2015), *Krt8-Cre^ERT2^* (Van Keymeulen et al., 2011), *Hi1fa^flox/flox^* (Garayoa et al., 2000), and *Il1r1^flox/flox^* (Robson et al., 2016) mice have been described and are available from Jackson Laboratory. *Il1r1-P2A-eGFP-IRES-CreERT2* (*Il1r1-Cre^ERT2^*) mice were generated in our laboratory. Mice for the lineage tracing and injury experiments were on a C57BL/6 background and 6-10 weeks old mice were used for most of the experiments described in this study. Experiments were approved by local ethical review committees and conducted according to UK Home Office project license PC7F8AE82. Mice were bred and maintained under specific-pathogen-free conditions at the Cambridge Stem Cell Institute and Gurdon Institute of University of Cambridge.

### Tamoxifen

Tamoxifen (Sigma) was dissolved in Mazola corn oil (Sigma) in a 20mg/ml stock solution. 0.2mg/g body weight tamoxifen was given via intraperitoneal (IP) injection. The numbers and date of treatment are indicated in the individual figures of experimental scheme.

### Bleomycin Administration

6-10 week mice were anesthetised via inhalation of isoflurane for approximately 3 mins. The mice were positioned on the intratracheal intubation stand, and 1.25U/kg of bleomycin, or PBS control, were delivered intratracheally by a catheter (22G). During the procedure anaesthesia was maintained by isoflurane and oxygen delivery.

### 3D Lung organoid co-culture

Lung organoids were established following the previous report (Lee et al., 2014). Briefly, freshly sorted lineage-labeled cells were resuspended in 3D basic media (DMEM/F12 (Gibco) supplemented with 10% FBS. (Gibco) and ITS (Insulin-Transferrin-Selenium, Corning)), and mixed with cultured lung stromal cells, followed by resuspension in growth factor-reduced Matrigel (BD Biosciences) at a ratio of 1:5. A 100 μl mixture was placed in a 24-well Transwell insert with a 0.4-μm pore (Corning). Approximately 5× 10^3^ *SPC*^+^ cells were seeded in each insert. 500 μL of 3D basic media was placed in the lower chamber, and medium was changed every other day with or without IL-1β (20ng/ml, Peprotech), Digoxin (50μM, Sigma), and 2-deoxyglucose (5mM, Sigma). ROCK inhibitor Y27632 (10uM, Sigma) was added in the medium for the first 2 days of culture. For isolation of stroma cells, cells negatively isolated by CD31 via MACS column were further negatively sorted by CD326 (EpCAM) and CD45 microbeads (Miltenyi Biotech). For co-culture with macrophages, sorted interstitial or alveolar macrophages were added to organoids with lineage-labeled *SPC*^+^ cells at a ratio of 1:6 in the presence of lung stromal cells. GM-CSF (20ng/ml, Peprotech) was included in some cultures. Analysis of colony forming efficiency (C.F.U) and size of organoids were at 14 days after plating if there is no specific description. For organoid culture of DATPs, AT2 cells (CD31^−^CD45^−^EpCAM^+^MHCII^+^) isolated from *Krt8-Cre^ERT2^;R26R^tdTomato^* were cultured with for 14 days with IL-1β (20ng/ml, Peprotech). Then, 4-OH tamoxifen was added at day14 and day16 in culture to label *Krt8*-expressing cells. Organoids were cultured with EpCAM^+^MHCII^−^Tomato^+^ DATPs isolated by flow cytometry.

### Lung tissue dissociation and flow cytometry

Lung tissues were dissociated with a collagenase/dispase solution as previously described. Briefly, after lungs were cleared by perfusion with cold PBS through the right ventricle, 2 mL of dispase (BD Biosciences, 50 U/ml) was instilled into the lungs through the trachea until the lungs inflated, followed by instillation of 1% low melting agarose (BioRad) through the trachea to prevent leakage of dispase. Each lobe was dissected and minced into small pieces in a conical tube containing 3 ml of PBS, 60 μL of collagenase/dispase (Roche), and 7.5 μL of 1% DNase I (Sigma) followed by rotating incubation for 45 min at 37°C. The cells were then filtered sequentially through 100-and 40-μm strainers and centrifuged at 1000rpm for 5 min at 4°C. The cell pellet was resuspended in 1ml of ACK lysis buffer (0.15 M NH4Cl, 10mM KHCO3, 0.1 mM EDTA) and lysed for 90 s at room temperature. 6 ml basic F12 media (GIBCO) was added and 500 μl of FBS (Hyclone) was slowly added in the bottom of tube. Cells were centrifuged at 1500 rpm for 5 min at 4°C. The cell pellet was resuspended in PF10 buffer (PBS with 10% FBS) for further staining. The antibodies used were as follows: CD45 (30-F11)-APC or-APC-Cy7 (BD Biosciences), CD31 (MEC13.3)-APC (BD Biosciences), Biotin-conjugated mouse lineage (Lin) panel that contains anti-B220 (RA3-6B2),-CD3ε(145-2C11),-Gr-1 (RB6-8C5),-CD11b (Mac-1, M1/70), -Ter-119 antibodies (Biolegend), EpCAM (G8.8)-PE-Cy7 or FITC (BioLegend), Sca-1 (Ly-6A/E, D7)–APC-Cy7 (BD Bioscience), MHC-II (I-A/I-E, M5)-FITC (eBiosceince), CD64 (X54-5/7.1)-PeCy7 (Biolegend), CD24(M1/69)-APC (eBioscience), and Siglec-F(E50-2440)-PE (BD Bioscience). 4’, 6-diamidino-2-phenylindole (DAPI) (Sigma) was used to eliminate dead cells. Data were acquired on LSRII analyzer (BD Biosceince) and then analyzed with FlowJo software (Tree Star). MOFLO system (Beckman Coulter) was used for the sorting at Wellcome-MRC Stem Cell Institute Flow Cytometry Facility.

### Human Adult Lung Tissue

Papworth Hospital NHS Foundation Trust (Research Tissue Bank Generic REC approval, Tissue Bank Project number T02233) provided deidentified lung samples obtained from IPF patients at the time of transplantation, normal background lung tissue from adult donor lungs that were deemed unsuitable for transplant, and lung adenocarcinoma tissues from lobectomies. Fresh tissues were fixed with 4% paraformaldehyde (PFA) overnight at 4°C and paraffin sections (7um) were used for immunofluorescent (IF) analysis.

### Macrophage culture *in vitro*

Interstitial macrophages (CD45^+^CD64^+^Siglec-F^−^CD11b^high^) or alveolar macrophages (CD45^+^CD64^+^Siglec-F^+^CD11^blow^) were isolated from C57BL/6 by MOFLO system (Beckman Coulter). Isolated macrophages were cultured for 24 hrs in RPMI-1640 medium containing 10% FBS and 50μM 2-mercaptoethanol with or without GM-CSF (10 ng/ml).

### EdU incorporation Assays in organoids

Lineage-labeled AT2 cells from *Il1r1*^flox/+^;R26R^tdTomato^ or *Il1r1*^flox/flox^*;R26R^tdTomato^* mice given by two doses of tamoxifen were isolated at day 4 post final injection. Organoids established in 8 well chamber slides (μ-Slide 8 wells, ibidi) were treated with EdU (10μM) at day 4 for 4 hrs. EdU staining was performed according to manufacturer’s instructions (Click-iT^®^ EdU Imaging Kits, Thermo Fisher Scientific).

### Measurement of Extracellular Acidification Rate (ECAR)

ECAR of organoids was measured using a XF94 analyzer (Seahorse Bioscience). Seahorse plates were pre-coated with 10% Matrigel in PBS for 1hr at 37°C. Organoids treated with PBS control or IL-1β were added with dispase to remove Matrigel and washed twice with XF Base Medium (DME, pH 7.4) supplemented with 1mM glutamine (Seahorse Bioscience). 30,000 cells were seeded on each well and incubated for 1hr at 37°C in non-CO_2_ incubator before measurement. Three components were injected automatically during the assay to achieve the following final concentrations: Glucose (10mM), Oligomycin (1μM), and 2-Deoxy Glucose (2-DG, 50mM). ECAR were normalized to the cell numbers of each wells.

### Glucose Uptake (2-NDBG incorporation) assays

Organoids at day 14 were washed twice with PBS and incubated with glucose-free medium supplemented with 10% FBS and GlutaMax (Gibco) for 1hr. 200μM of 2-NBDG (Life Technologies) were subsequently added for 1hr. Organoids were dissociated into single cells with trypLE Express (Gibco) and cells were harvested for flow cytometry. A control sample lacking 2-NBDG was used to set the flow cytometer compensation and gate parameters for 2-NBDG positive events.

### Quantitative RT-PCR

Total RNA was isolated using TRI-reagent (Molecular Research Center) or using a Qiagen RNeasy Micro Kit according the manufacturer’s instructions. Equivalent quantities of total RNA were reverse-transcribed with SuperScript cDNA synthesis kit (Life Technology) or QuantiTect (Qiagen). Diluted cDNA was analyzed by real-time PCR (StepOnePlus; Applied Biosystem). Pre-designed probe sets and TaqMan universal PCR Master Mix (2x, Thermo Fisher Scientific) were used as follows: Ager (Mm_00545815_m1), Pdpn (Mm_00494716_m1), Aqp5 (Mm_00437578_m1). Gapdh expression (Mm_00805216_m1) was used to normalise samples using the ΔCt method. Sybr green assays were also used with SYBR Green Master Mix (2x, Thermo Fisher Scientific). Primer sequences are as follows:

Gapdh: F-AGGTCGGTGTGAACGGATTTG, R-TGTAGACCATGTAGTTGAGGTCA

Vegfa: F-CCGGTTTAAATCCTGGAGCG, R-TTTAACTCAAGCTGCCTCGC

Clic5: F-ATGACGGACTCAGCGACAAC, R-GTAGATCGGCTGGCTTTCTTTT

Cav-1: F-TGAGAAGCAAGTGTATGACGC, R-CTTCCAGATGCCGTCGAAAC

Aqp5: F-TCTTGTGGGGATCTACTTCACC, R-TGAGAGGGGCTGAACCGAT

Sdpr: F-GCTGCACAGGCAGAAAAGTTC, R-GTGACAGCATTCACCTGCG

Spock2: F-ACCCCCGGCAATTTCATGG, R-TGTCTTCCCAGCTCTTGATGTAA

Limch2: F-AAAGGCCCTTCAGATACGGTC, R-TACTCGTGCTCTCTGCGTCAT

Etv5: F-TCAGTCTGATAACTTGGTGCTTC, R-GGCTTCCTATCGTAGGCACAA

Abca3: F-CAGCTCACCCTCCTACTCTG, R-ACTGGATCTTCAAGCGAAGCC

Lpcat1: F-GGCTCCTGTTCGCTGCTTT, R-TTCACAGCTACACGGTGGAAG

Itga7: F-CTGCTGTGGAAGCTGGGATTC, R-CTCCTCCTTGAACTGCTGTCG

Lrg1: F-TTGGCAGCATCAAGGAAGC, R-CAGATGGACAGTGTCGGCA

Orm1: F-CGAGTACAGGCAGGCAATTCA, R-ACCTATTGTTTGAGACTCCCGA

Slc2a1: F-CAGTTCGGCTATAACACTGGTG, R-GCCCCCGACAGAGAAGATG

Slc16a3: F-TCACGGGTTTCTCCTACGC, R-GCCAAAGCGGTTCACACAC

Cldn4: F-GTCCTGGGAATCTCCTTGGC, R-TCTGTGCCGTGACGATGTTG

Hif1a: F-ACCTTCATCGGAAACTCCAAAG, R-ACTGTTAGGCTCAGGTGAACT

IL-1β: F-GCAACTGTTCCTGAACTCAACT, R-ATCTTTTGGGGTCCGTCAACT

IL-13: F-CCTGGGCTCTTGTCTGCCTT, R-GGTCTTGTTGATGTTGCTCA

IL-18: F-GACTCTTGCGTCAACTTCAAGG, R-CAGGCTGTCTTTTGTCAACGA

IL-22: F-ATGAGTTTTTCCCTTATGGGGAC, R-GCTGGAAGTTGGACACCTCAA

IL-33: F-TCCAACTCCAAGATTTCCCCG, R-CATGCAGTAGACATGGCAGAA

Fgf7: F-TTTGGAAAGAGCGACGACTT, R-GGCAGGATCCGTGTCAGTAT

IL-6: F-TCTATACCACTTCACAAGTCGGA, R-GAATTGCCATTGCACAACTCTTT

### Histology and Immunohistochemistry

Mouse lung tissues were routinely perfused, inflated, and fixed with 4% PFA for 4-6 hrs at 4 degrees and cryosections (8um) and paraffin sections (7um) were used for histology and IF analysis. Cultured colonies from organoids were fixed with 4% PFA for 2-4 hrs at room temperature followed by immobilization with Histogel (Thermo Scientific) for paraffin embedding. Sectioned lung tissues or colonies were stained with hematoxylin and eosin (H&E) or immunostained: after antigen retrieval with citric acid (0.01M, pH 6.0), blocking was performed with 5% normal donkey serum in 0.2% Triton-X/PBS at room temperature for 1hr. Primary antibodies were incubated overnight at 4°C at the indicated dilutions: goat anti-SP-C (1:200, Santa Cruz Biotechnology Inc., sc-7706), pro-SP-C (1:300, Millipore, AB3786), rabbit anti-Ki67 (1:250, A. Menarini, MP-325-CRM1), rat anti-Ki67 (1:200, Biolegend, A16A8), rabbit anti-RFP (1:250, Rockland, 600–401379), hamster anti-PDPN (1:1000, DSHB, 8.1.1), rat anti-Cytokeratin-8 (1:300, DSHB, TROMA-I), rabbit anti-Claudin-4 (1:200, Thermo Fisher Scientific, 36-4800), rabbit anti-Hopx (1:100, Santa Cruz Biotechnology Inc., sc-30216), rabbit anti-Aqp5 (1:200, Alomone Labs, AQP5-005), rabbit anti-Caveolin-1 (1:500, Cell Signaling, #3267), and mouse anti-HTII-280 (1:200, Terrace Biotechnology, TB-27AHT2-280). Alexa Fluor-coupled secondary antibodies (1:500, Invitrogen) were incubated at room temperature for 60 min. After antibody staining, nuclei were stained with DAPI (1:1000, Sigma) and sections were embedded in Vectashield (Vector Labs). Fluorescence images were acquired using a confocal microscope (Leica TCS SP5). All the images were further processed with Fiji software.

### Statistical Analysis

Statistical methods relevant to each figure are outlined in the figure legend. Statistical analyzes were performed with Prism software package version 7.0 (GraphPad). *P* values were calculated using two-tailed unpaired or paired Student’s t test. Sample size for animal experiments was determined based upon pilot experiments. Mice cohort size was designed to be sufficient to enable accurate determination of statistical significance. No animals were excluded from the statistical analysis. Mice were randomly assigned to treatment or control groups, while ensuring inclusion criteria based on gender and age. Animal studies were not performed in a blinded fashion. The number of animals shown in each figure is indicated in the legends as *n* = *x* mice per group. Data shown are either representative of three or more independent experiments, or combined from three or more independent experiments as noted and analyzed as mean ± SEM.

### Cell Counting and Image Analysis

Sections included in cell scoring analysis for lung tissue were acquired using Leica TCS SP5 confocal microscope. At least five different sections including at least 10 alveolar regions from three individual mice per group were used. Cell counts were performed on ImageJ using the ‘Cell Counter’ plug-in and the performer was blinded to the specimen genotype and condition. At least two step sections (30um apart) per individual well were used for quantification of AT1 or AT2 cells.

### ATAC-seq analysis

The ATAC-seq assay was performed on 50,000 FACS-purified cells as previously described (Buenrostro et al., 2015). In brief, two biological independent samples were used for ATAC-seq experiment. 5 mice were pooled for *Il1r1*^+^AT2 cells and 1 mouse was used for bulk AT2 cells per group. Purified cells were lysed in ATAC lysis buffer for 5 min to get nuclei and then transposed with Tn5 transposase (Illumina) for 30 min. Fractionated DNA was used for amplification and library preparation according to manufacturer’s guidelines (Illumina) and 150 bp-paired end sequencing was performed by pooling two samples of *Il1r1*^+^AT2 and bulk AT2 cells, respectively, in one lane of the Illumina HiSeq 4000 platform. The quality of the generated sequencing data was checked using the FastQC program, followed by filtering of adaptor and/or overrepresented sequences using Trimmomatic (Bolger et al., 2014). Filtered reads were next mapped to the mouse primary genome assembly (mm9/GRCm38) using STAR (Dobin et al., 2013), with parameters – outFilterMatchNminOverLread 0.4–outFilterScoreMinOverLread 0.4, and a GTF annotation file of the latest mouse assembly (GCA_000001635.8) downloaded from ENSEMBL ftp. Duplicate reads were flagged and removed using MarkDuplicates from Picard tools. MACS2(Feng et al., 2012) callpeak was used for ATAC-seq peak calling of the *Il1r1*^+^AT2 and bulk AT2 samples, using the options–nomodel–shift-100–extsize 200. Differentially enriched peaks in *Il1r1*^+^AT2 and bulk AT2 populations were next inferred using the MACS2 bdgdiff with a log10 likelihood ratio score cutoff of 10. ATAC-seq heatmaps were plotted using deepTools2 (Ramirez et al., 2016). Annotation of ATAC-seq enriched peaks overlapping with promoter and other gene regions was performed using the ChIPseeker R/Bioconductor package, together with GO enrichment and pathway analyzes (Yu et al., 2015). Finally, motif identification was performed using the findMotifsGenome.pl program of the HOMER software (Heinz et al., 2010).

### scRNA-seq Library Preparation and Sequencing

Established organoids of control or IL-1β-treatment were incubated with dispase (BD Bioscience) for 30-60min. Then, cells were dissociated with TripLE (Gibco) for 5min, followed by washing with buffer (PBS/0.01% BSA). For *SPC lineage-*labeled cells, CD45^−^CD31^−^EpCAM^+^Tomato^+^ cells were sorted at specific time points (at day 14 and day 28 post damage) from PBS or Bleomycin-treated mice (2 mice were pooled for each experiment). For non-lineage-labeled cells isolated from *SPC-Cre^ERT2^;R26R^tdTomato^* mice in parallel with experiment of *SPC* lineage-labeled cells, we combined the cells of EpCAM^+^Tomato^−^ and EpCAM^−^ population with a ratio of 2:1, respectively. The resulting cell suspension (~110,000 cells each) were submitted as separate samples to be barcoded for the droplet-encapsulation single-cell RNA-seq experiments using the Chromium Controller (10X Genomics). Single cell cDNA synthesis, amplification and sequencing libraries were generated using the Single Cell 3’ Reagent Kit as per the 10x Genomics protocol. Libraries were multiplexed so that 2 libraries were sequenced per single lane of HiSeq 4000 using the following parameters: Read1: 26 cycles, i7: 8 cycles, i5: 0 cycles; Read2: 98 cycles to generate 75bp paired end reads.

### Alignment, quantification and quality control of single cell RNA sequencing data

Droplet-based sequencing data was aligned and quantified using the Cell Ranger Single-Cell Software Suite (version 2.0.2, 10x Genomics Inc) using the *Mus musculus* genome (GRCm38) (official Cell Ranger reference, version 1.2.0). Cells were filtered by custom cutoff (more than 500 and less than 7000 detected genes, more than 2000 UMI count) to remove potential empty droplets and doublets. Downstream analysis included data normalisation, highly variable gene detection, log transformation, principal component analysis, neighbourhood graph generation and Louvain graph-based clustering, which was done by python package scanpy (version 1.3.6) (Wolf et al., 2018) using default parameters.

### Excluding stromal cells and contaminated cells in scRNA-seq analysis of organoids and *SPC* lineage-tracing after bleomycin injury

For scRNA-seq analysis of organoids, we excluded the cluster of EpCAM^−^ cells of stromal cells we put together with AT2 cells in culture. For *in vivo* scRNA-seq analysis of AT2 cells after bleomycin injury, we excluded non-epithelial cells and ciliated cells based on marker gene expression. Although cells were sorted based on the expression of EpCAM, CD31, CD45, and Tomato before scRNA-seq, 255 contaminating cells among 12514 cells captured were observed in the initial droplet dataset. These comprised: 214 ciliated cells expressing *Foxj1*, *Wnt7b*, and *Cd24a*; 16 mesenchyme cells expressing *Vcam1*, *Acta2*, *Des*, and *Pdgfra*; 25 immune cells expressing *Ptprc* (CD45), *Tyrobp*, *Il2rg*, and *Lck*. Each of these cell populations was identified by an initial round of unsupervised Louvain graph-based clustering analysis as they formed extremely distinct clusters and then removed. For scRNA-seq analysis of *in vivo* non-lineage-labeled cells, we excluded the doublet cluster of cells expressing both EpCAM^+^CD45^+^ (1125 cells among 14017 cells).

### Doublet Exclusion

To exclude doublets from single-cell RNA sequencing data, we applied scrublet algorithm per sample to calculate scrublet-predicted doublet score per cell with following parameters: sim_doublet_ratio = 2; n_neighbors=30; expected_doublet_rate= 0.1. Any cell with scrublet score > 0.7 was flagged as doublet. To propagate the doublet detection into potential false-negatives from scrublet analysis, we over-clustered the dataset (*sc.tl.louvain* function from scanpy package version 1.3.4; resolution = 20), and calculated the average doublet score within each cluster. Any cluster with averaged scrublet score > 0.6 was flagged as a doublet cluster. All remaining cell clusters were further examined to detect potential false-negatives from scrublet analysis according to the following criteria: (1) Expression of marker genes from two distinct cell types which are unlikely according to prior knowledge, (2) higher number of UMI counts.

### Pseudotime Analysis

All data contained within our processed Seurat object for the wildtype data set was converted to the AnnaData format for pseudotime analysis in Scanpy (version 1.3.6). We recalculated *k*-nearest neighbors at k = 15. Pseudotime was calculated using Scanpy’s partitioned-based graph abstraction function, PAGA. Diffusion pseudotime was performed using Scanpy’s DPT function with default parameters.

## Supplemental Information

**Figure S1, related to Fig. 1.**
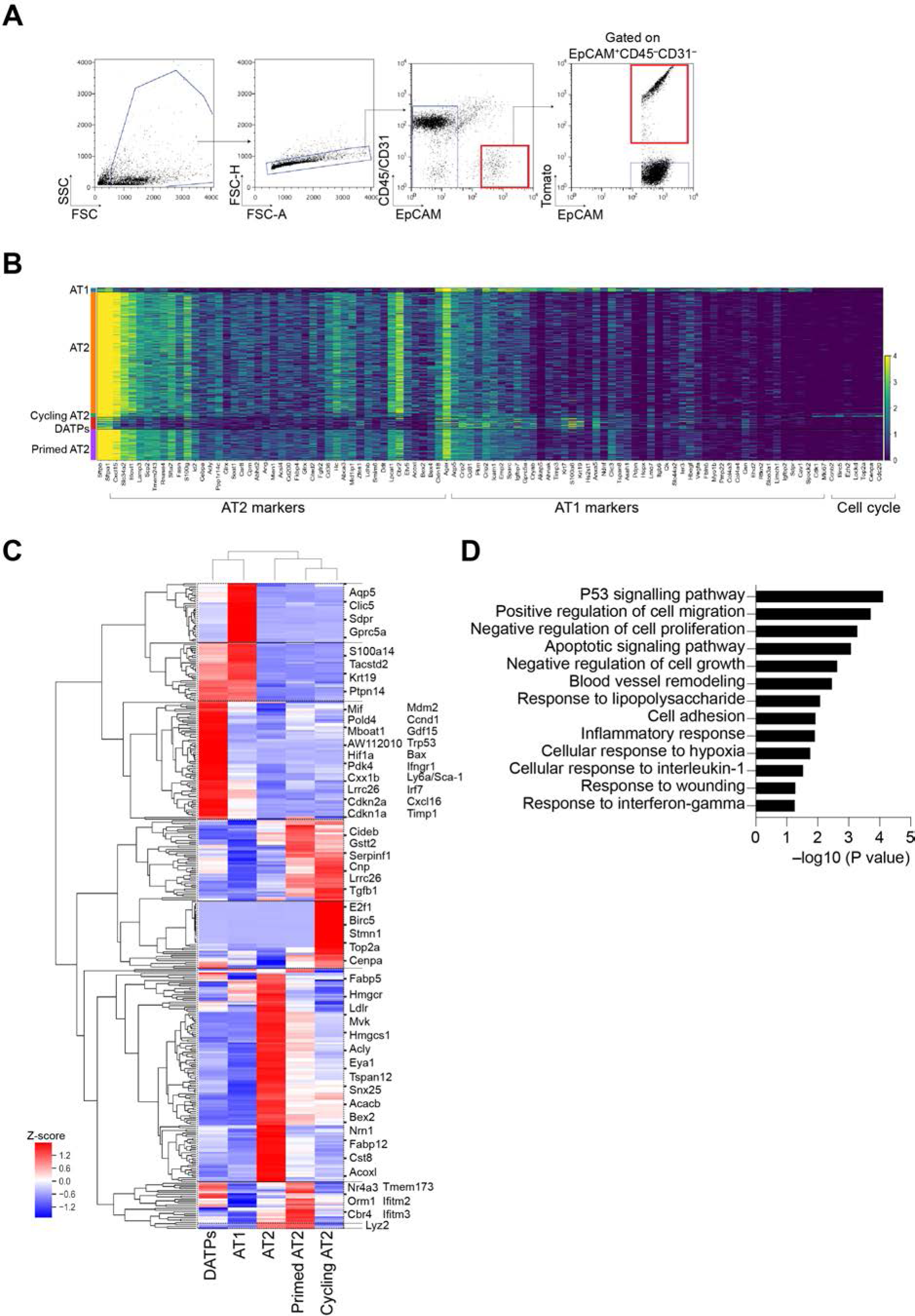
Single-cell profiling of *SPC* lineage-labeled cells during injury repair. **(A)** Sorting strategy for *SPC* lineage-labeled cells by flow cytometry after bleomycin injury. **(B)** Gene expression of AT2 markers, AT1 markers, or cell cycle markers across single cells from distinctive subsets revealed by single-cell RNA sequencing (scRNA-seq) analysis during injury repair. **(C)** Heap map showing relative expression of marker genes in distinctive subsets revealed by scRNA-seq analysis. **(D)** GO analysis of enriched genes in DATPs.

**Figure S2, related to Fig. 2.**
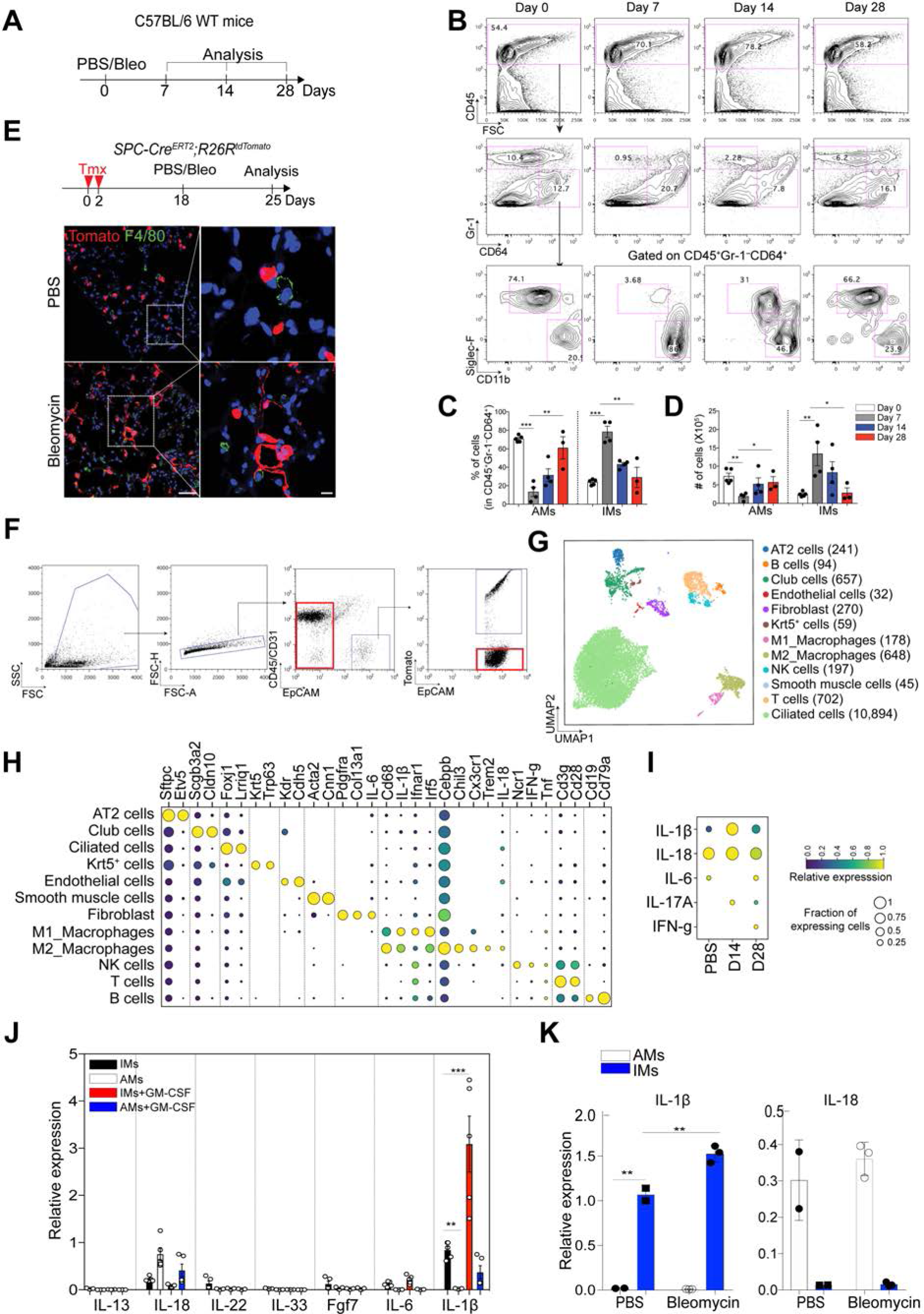
Dynamics of macrophages during alveolar regeneration after bleomycin injury. **(A)** Schematic of experimental design for analysis of immune cells at indicated time points after bleomycin injury. **(B)** Flow cytometry analysis of alveolar (Siglec-F^+^CD11^blow^) and interstitial (Siglec-F^−^ CD11b^high^) macrophages at indicated time points post injury. Cells gated on CD45^+^CD64^+^Gr-1^−^ were analyzed further for expression of Siglec-F and CD11b. Numbers adjacent to the outlined area indicate the percentage of populations. **(C, D)** Frequencies (**C**) and absolute cell numbers (**D**) of alveolar (AMs) or interstitial (IMs) macrophages at indicated time points. Each individual dot represents one experiment and date are presented as mean ± SEM. *p<0.05, **p<0.01, and ***p<0.001. **(E)** Experimental design (top) of *SPC* lineage-tracing and immunofluorescent (IF, bottom) images of tissue samples after bleomycin treatment. IF images show the increased numbers of F4/80+ macrophages at day 7 post injury. A high magnification images (right) show the interaction between macrophages and *SPC* lineage-labeled cells. Data are the representative of two independent experiments. Scale bar, 50 μm (left) and 10 μm (right). Tomato (red), F4/80 (green), and DAPI (blue). **(F)** Sorting strategy for *SPC* unlabeled single cells pooling of EpCAM^+^Tomato^−^ and EpCAM^−^ population by flow cytometry after bleomycin injury. **(G)** Clusters of unlabeled cells (14,017) after bleomycin injury from 10xGenomics 3’ scRNA-seq analysis visualized by UMAP, assigned by specific colors. Number of cells in the individual cluster is depicted in the figure. **(H)** Gene expression of key markers in each distinctive cluster. *IL-1β* is specifically expressed in macrophages. **(I)** Gene expression of *IL-1β*, *IL-18*, *IL-6*, *IL-17A*, and *IFN-g* at indicated time points after bleomycin injury. Of note, the expression of *IL-1β* is dramatically increased at day 14 post injury and returns back to the homeostatic level at day 28 post injury. **(J)** qPCR analysis of specific cytokine expression in alveolar (AMs) or interstitial (IMs) macrophages in response to activation by GM-CSF. Isolated subsets of macrophages were cultured in the presence or absence of GM-CSF for 24hrs *in vitro*. Each individual dot represents one experiment and date are presented as mean ± SEM. **(K)** qPCR analysis for *IL-18* and *IL-1β* in alveolar (AMs, white bar) or interstitial (IMs, blue bar) macrophages isolated at day 7 after PBS or bleomycin treatment. Each individual dot represents one experiment from one mouse and date are presented as mean ± SEM. **p<0.01, ***p<0.001.

**Figure S3, related to Fig. 2, F-H.**
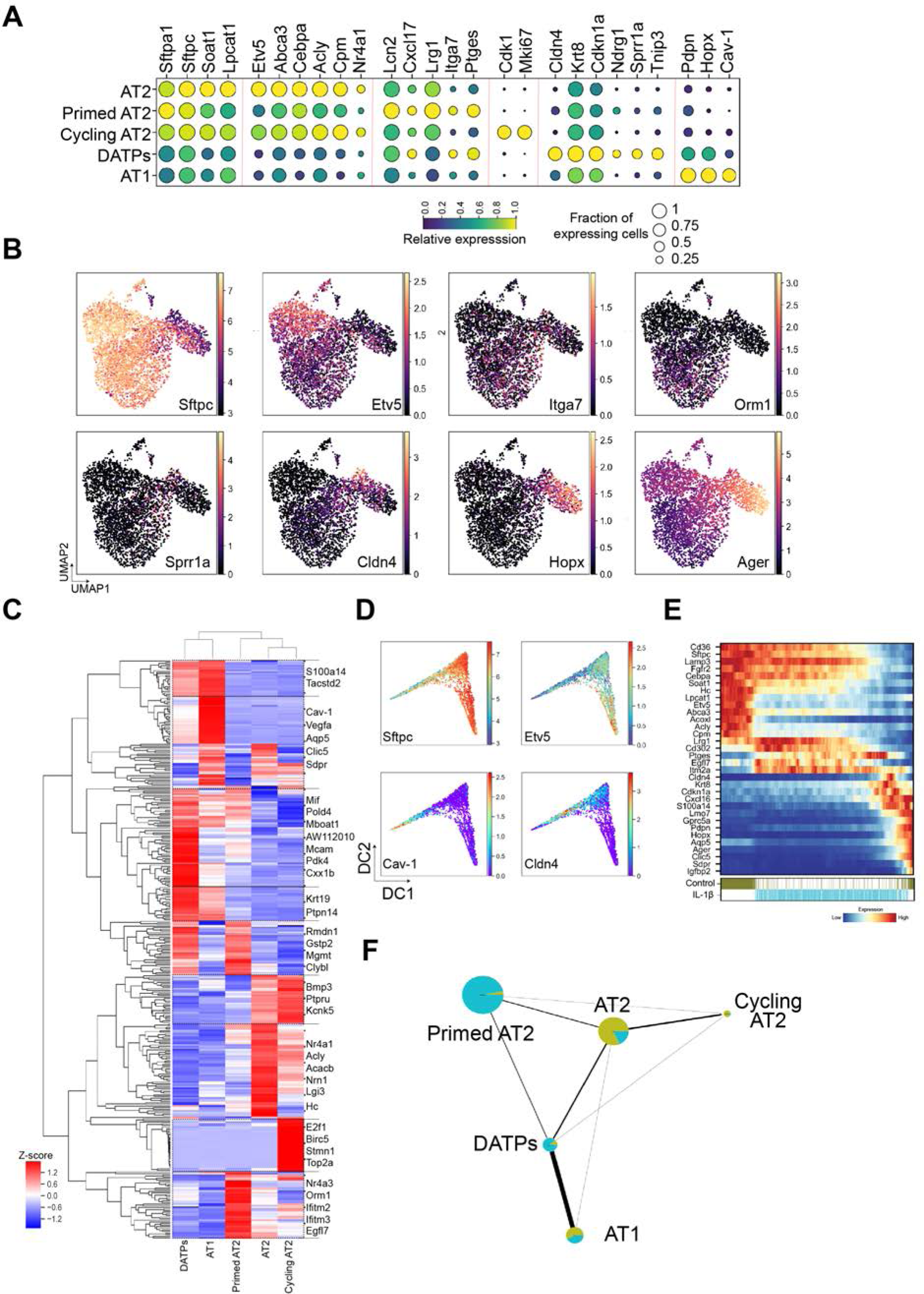
Alveolar organoids challenged by IL-1β recapitulate the behavior of regenerating AT2 cells during injury repair. **(A)** Gene expression of key markers in each distinctive cluster. **(B)** UMAP visualization of the log-transformed (log_10_(TPM+1)), normalized expression of selected marker genes in distinctive clusters. **(C)** Heap map showing relative expression of selected genes that are specifically expressed in distinctive clusters revealed by scRNA-seq analysis. **(D)** Diffusion map according to diffusion pseudotime order colored by expression (log_10_(TPM+1)) of specific genes. **(E)** Gene expression profiles of control and IL-1β-treated organoids ordered according to pseudotime trajectory. Lower color bars indicate annotation by samples. **(F)** Network topology among clusters from single cell data revealed by Partition-based graph abstraction (PAGA). Colors indicate the proportion of each cluster by time point. Each node in the PAGA graph represents a cluster and the weight of the lines represents the statistical measure of connectivity between clusters.

**Figure S4, related to Fig. 3.**
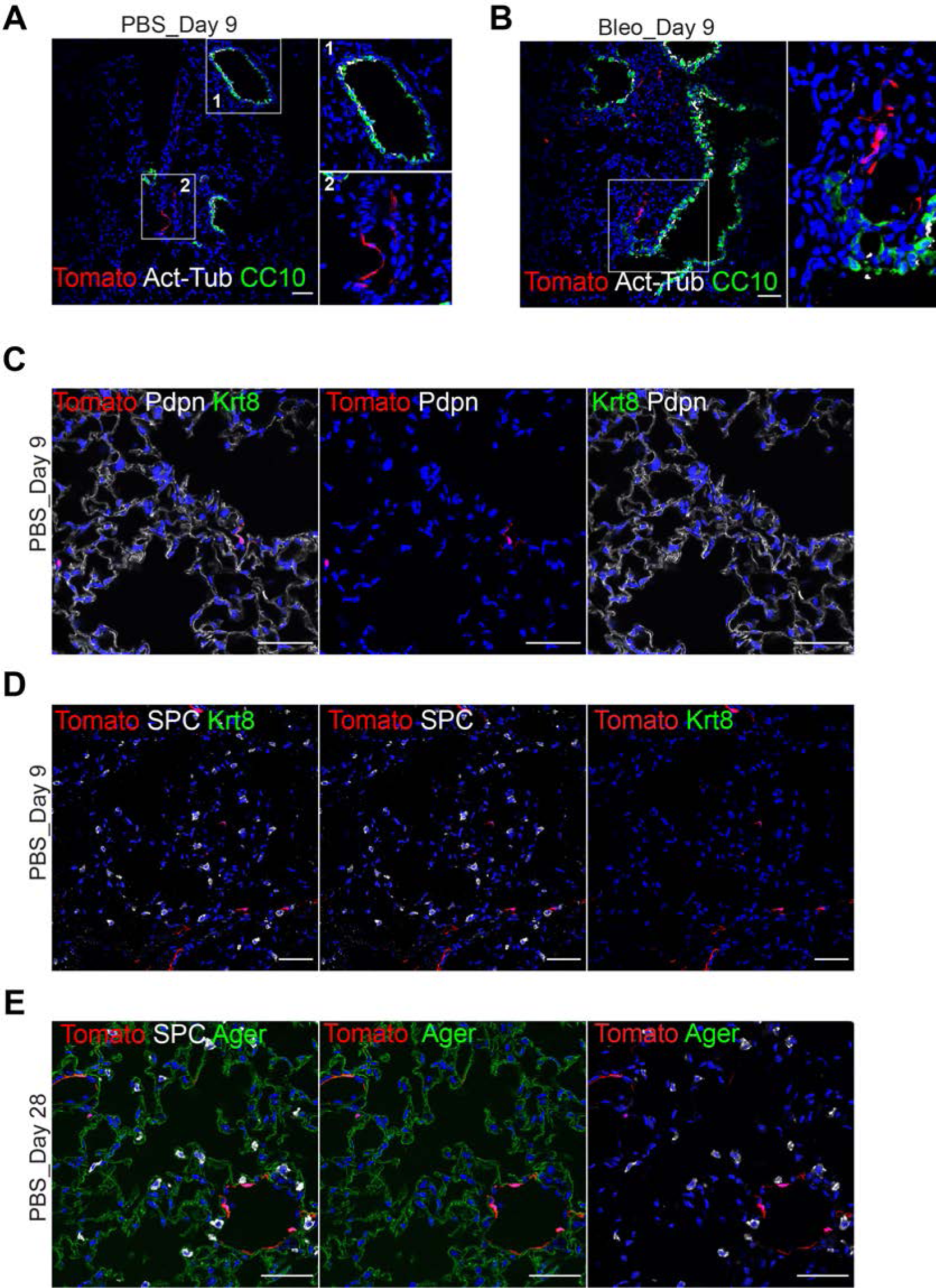
*Ndrg1* does not mark epithelial cells in steady-state lungs. **(A)** Representative IF images show that airway cells are not marked by *Ndrg1* expression at day 9 post PBS treatment: Tomato (for *Ndrg1* lineage, red), CC10 (green, club cells), Acetyl-Tub (white, ciliated cells), and DAPI (blue). Insets (left) show high-power view (right). **(B)** Representative IF images show that airway cells are not marked by *Ndrg1* expression at day 9 post bleomycin injury: Tomato (for *Ndrg1* lineage, red), CC10 (green, club cells), Acetyl-Tub (white, ciliated cells), and DAPI (blue). Insets (left) show high-power view (right). **(C)** Representative IF images show that lineage-labeled DATPs are barely observed at day 9 post PBS treatment: Tomato (for *Ndrg1* lineage, red), Pdpn (white) and Krt8 (green). Scale bar, 50 μm. **(D)** Representative IF images show that SPC+ AT2 cells are not labeled by *Ndrg1* expression at day 9 post PBS treatment: Tomato (for *Ndrg1* lineage, red), SPC (white) and Krt8 (green). Scale bar, 50 μm. **(E)** Representative IF images show that neither AT2 and AT1 cells are labeled by *Ndrg1* expression at day 28 post PBS treatment: Tomato (for *Ndrg1* lineage, red), SPC (white) and Ager (green). Scale bar, 50 μm.

**Figure S5, related to Fig. 3.**
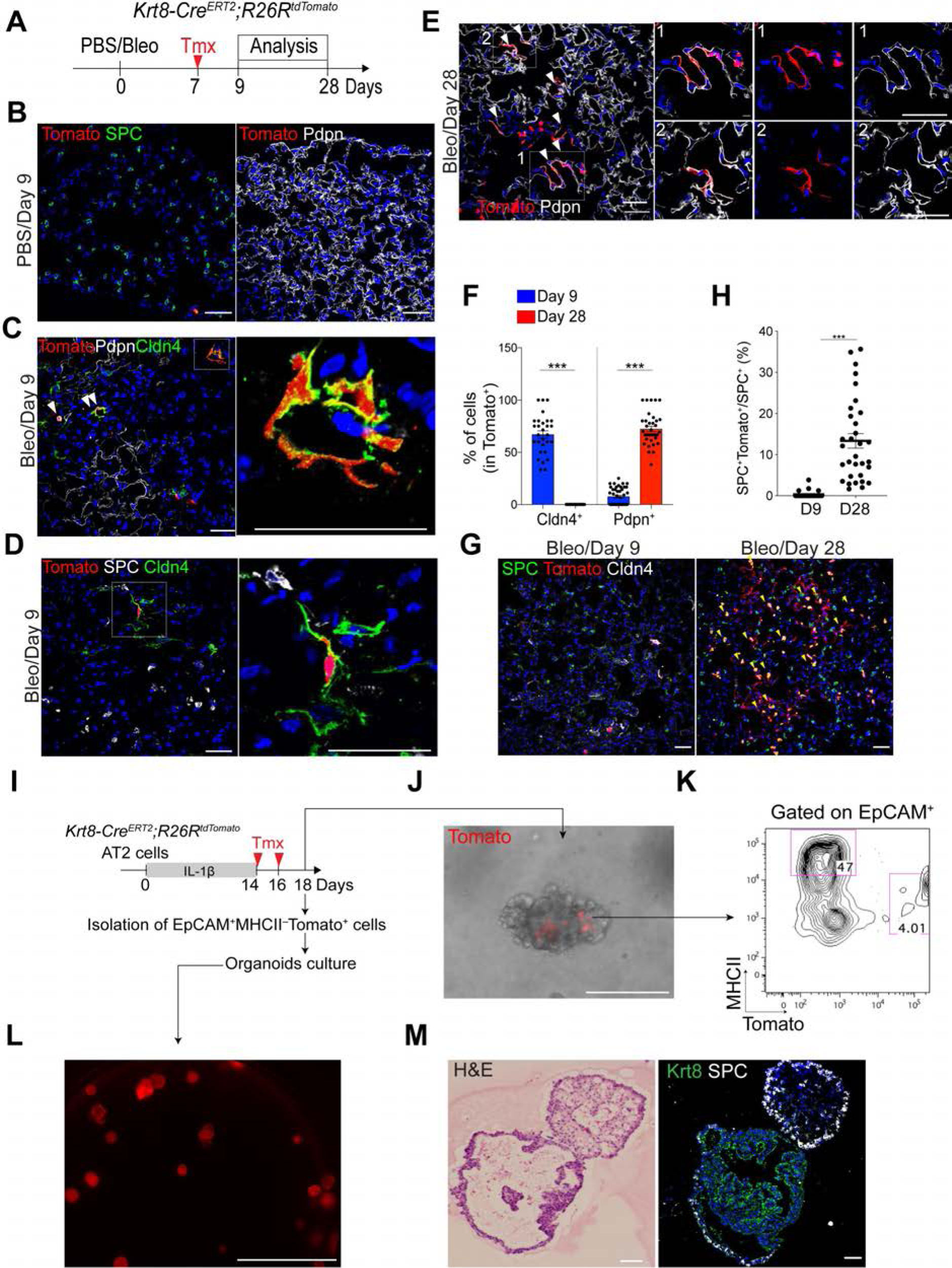
Lineage tracing analysis of *Krt8*^+^ cells reveals that DATPs are capable of producing AT1 cells and reverting to AT2 cells during alveolar regeneration. **(A)** Experimental design for *Krt8* lineage-tracing analysis using *Krt8-Cre^ERT2^;R26R^tdTomato^* reporter mice after bleomycin injury. Specific time points for tamoxifen injection and analysis are indicated. **(B)** Representative IF images show that none of AT2 (left) and AT1 (right) cells are lineage-labeled by *Krt8* expression in uninjured lung (PBS control): Tomato (red), SPC (green, left), Pdpn (white, right), and DAPI (blue). Scale bar, 50 μm. **(C, D)** Representative IF images show that *Krt8* lineage-labeled cells express Cldn4 at day 9 post injury. None of AT1 (**C**) and AT2 (**D**) cells are lineage-labeled by *Krt8* expression at this time point: Tomato (red), Pdpn (white), Cldn4 (green), and DAPI (blue). Arrowhead points to *Krt8* lineage-labeled DATPs. White boxed insets are shown on the right. Scale bar, 50 μm. **(E)** Representative IF images show that *Krt8* lineage-labeled cells generate new AT1 cells at day 28 after injury: Tomato (red), Pdpn (white), and DAPI (blue). Arrowhead points to lineage-labeled Pdpn^+^ cells. Insets (left) show high-power view (1, right top; 2, right bottom). Scale bar, 50 μm. **(F)** Statistical quantification of Cldn4^+^Tomato^+^ or Pdpn^+^Tomato^+^ cells at indicated time points after injury. Each individual dot represents one section and data are presented as mean ± SEM with two independent experiments (n=5). ***p<0.001. **(G)** Representative IF images show that *Krt8* lineage-labeled cells generate AT2 cells at day 28 post injury. Notably, there are few AT2 cells that are marked by *Krt8* expression at day 9 post injury: Tomato (for *Krt8* lineage, red), SPC (green), Cldn4 (white), and DAPI (blue). Arrowhead points to lineage-labeled AT2 cells. Scale bars, 50 μm. **(H)** Quantification of *Krt8* lineage-labeled SPC^+^ AT2 cells. Each individual dot represents one section and data are presented as mean ± SEM with three independent experiments (n=4). ***p<0.001. **(I)** Scheme of experimental design for organoid culture assays. AT2 cells were isolated by surface markers CD31^−^CD45^−^EpCAM^+^MHCII^+^ from *Krt8-Cre^ERT2^;R26R^tdTomato^* mice and cultured as organoids with IL-1β for 14 days. 4-OH tamoxifen was added at day14 and day16 in culture to label *Krt8*-expressing cells. At day 18, organoids were further analyzed for a microscopy (I), flow cytometry (J), and organoid formation (K and L). **(J)** Representative merged fluorescent and brightfield image of organoids in (H). Treatment of 4-OH tamoxifen allows to mark *Krt8*^+^ (Tomato^+^) cells. Scale bar, 200 μm. Notably, Tomato signals were detected only in inner parts of organoids. **(K)** Flow cytometry analysis of AT2 (EpCAM^+^MHCII^+^Tomato^−^) and DATPs (EpCAM^+^MHCII^−^Tomato^+^) from dissociated organoids in (I). Numbers adjacent to the outlined area indicate the percentage of populations. Of note, Tomato^+^ cells are not AT2 cells. **(L, M)** Representative fluorescent image (**K**), and H&E staining (**L**, left) and IF image (**L**, right) of organoids derived from dissociated *Krt8*^+^Tomato^+^ cells in (I and J). Scale bar, 1,000 μm (**K**) and 50 μm (**L**).

**Figure S6, related to Fig. 4, B-E.**
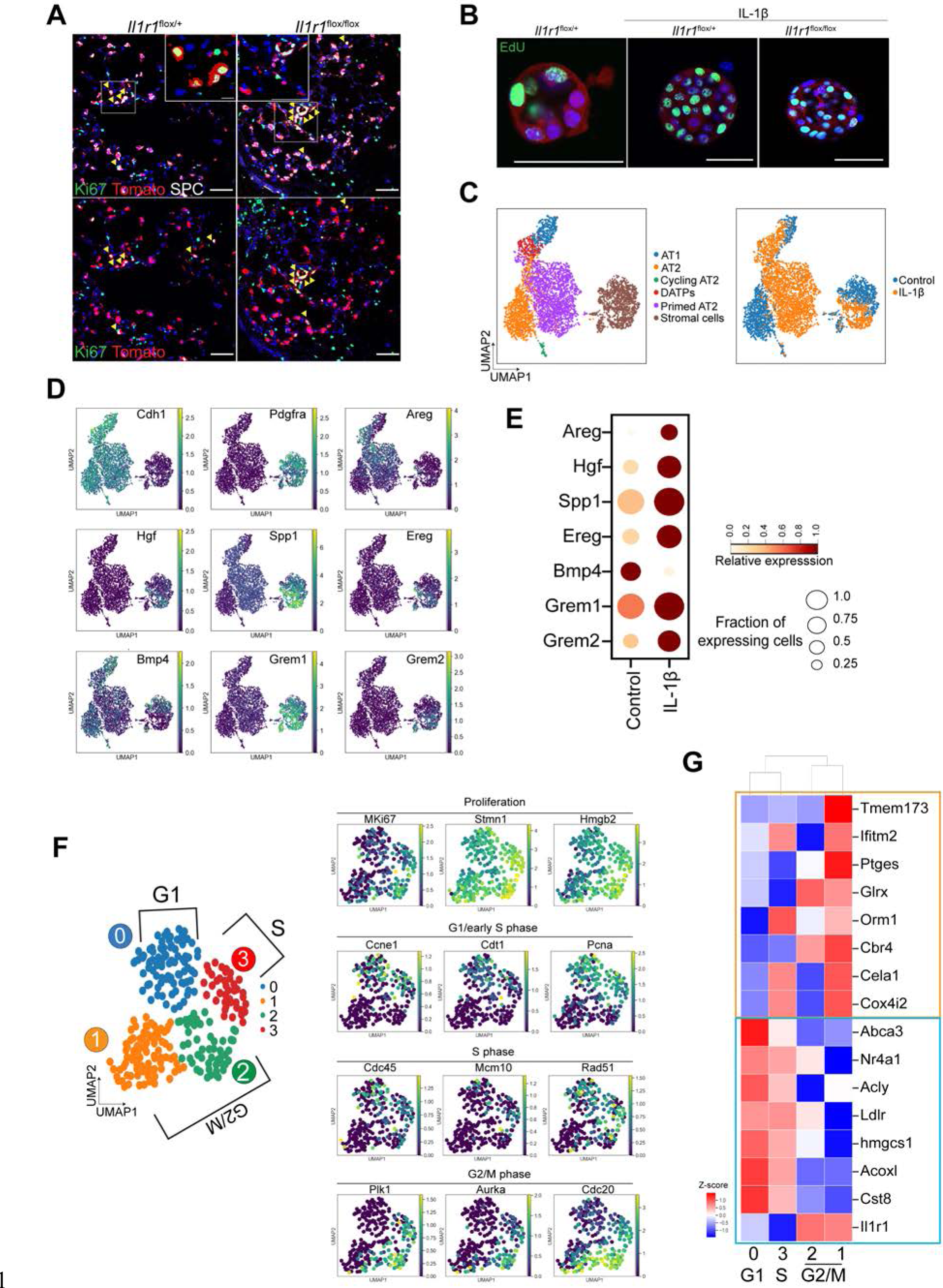
IL-1β signaling primes AT2 cells during cell cycle transition. **(A)** Representative IF images showing Ki67^+^ lineage-labeled AT2 cells in the lung of mice treated with PBS or bleomycin at day 7 post injury: Tomato (for *SPC* lineage, red), Ki67 (green), SPC (white), and DAPI (blue). Arrowheads, Ki67^+^ AT2 cells. Insets show high-power view. Scale bars, 50 μm. No discernible differences in number of Ki67^+^ AT2 cells were observed in the lung of indicated genotyped mice. **(B)** Representative IF images showing proliferating cells in AT2 organoids derived from the lungs of indicated genotyped mice. Organoids were pulsed with BrdU for 4hrs at day 4 in cultures. Notably, IL-1β treatment enhances proliferation in organoids regardless of *Il1r1* expression in AT2 cells. **(C)** UMAP visualization of cell clusters from scRNA-seq analysis of epithelial cells and stromal cells from control or IL-1β-treated organoids. Cells were isolated at day 21 in organoid culture. Colors indicate distinct cell types (left) and samples (right). **(D)** UMAP visualization of the log-transformed (log_10_(TPM+1)), normalized expression of cell type marker genes (e.g. *Cdh1* for epithelial cells and *Pdgfra* for stromal cells/fibroblast) and growth factors in each distinctive cluster. **(E)** Gene expression of growth factors that may enhance proliferation of AT2 cells in control or IL-1β-treated stromal cells. **(F)** Clusters of Cycling AT2 population (cAT2) shown in Fig. 1B visualized by UMAP, assigned by specific colors. Based on the expression of cell cycle genes, four clusters were classified into two cell cycle phases; G1 (cluster 0), S phase (cluster 3) and G2/M phase (cluster 2 and 1). UMAP visualization of the log-transformed (log_10_(TPM+1)), normalized expression of marker genes for cell proliferation and cell cycle (G1/early S phase; S phase; G2/M phase). **(G)** Heap map showing the *Il1r1* expression and acquisition of Primed AT2 cell (pAT2) signatures during cell cycle transition. Acquisition of transcriptional signatures of pAT2 cells by downregulating of naïve AT2 cell markers including *Abca3* (blue box) and inducing expression of genes related with inflammatory response including *Ptges* (orange box) during cell cycle transition from S to G2/M phase.

**Figure S7, related to Fig. 4F.**
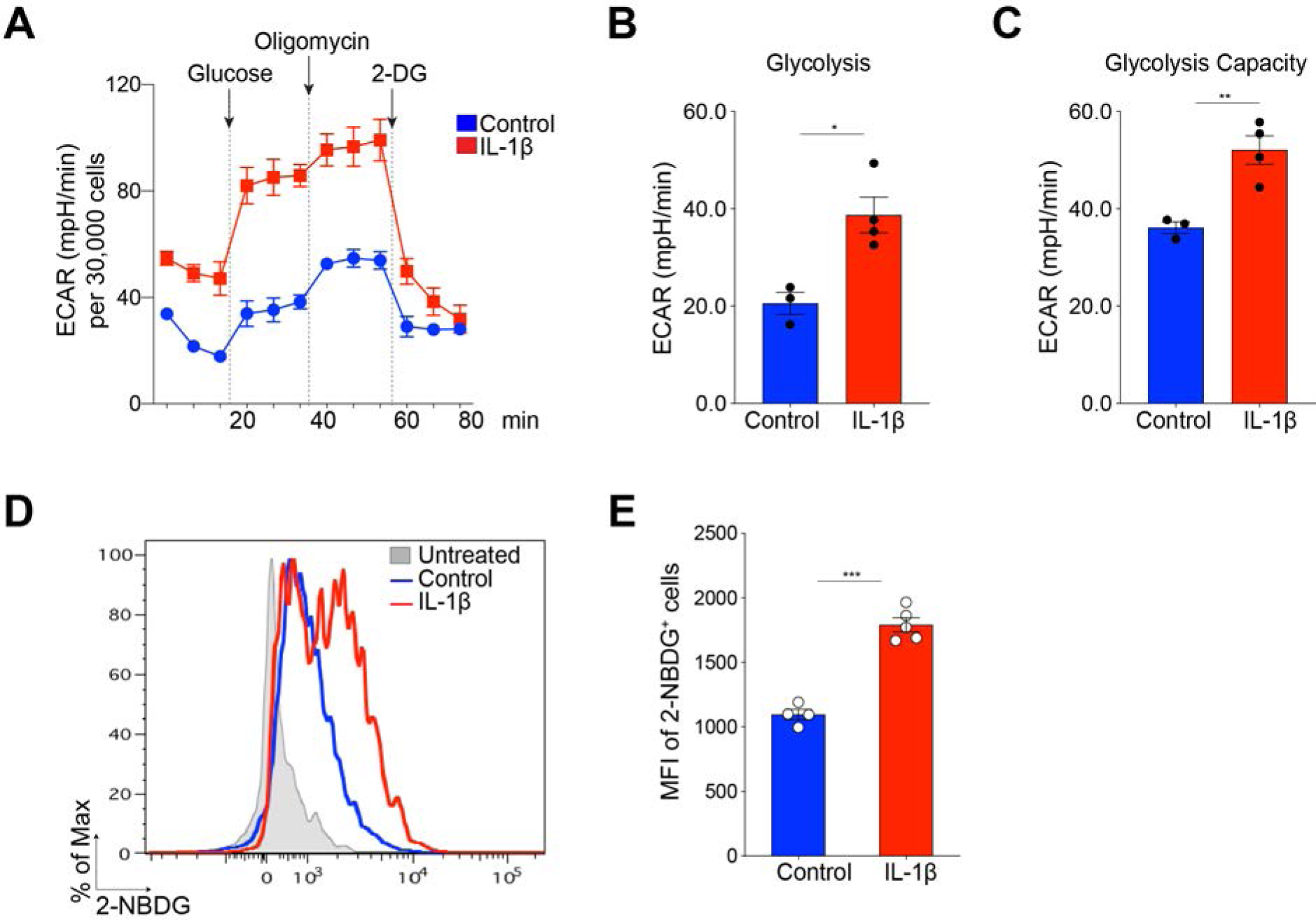
IL-1β enhances the glycolysis metabolism. **(A)** Real-time ECAR (<u>E</u>xtra<u>c</u>ellular <u>A</u>cidification <u>R</u>ate) of organoids treated with PBS (control) and IL-1β was measured by XF-96 analyzer. Vertical lines with arrow indicate addition of glucose (glycolysis substrate, 10mM), oligomycin (ATP synthase inhibitor, 1uM), and 2-Deoxy Glucose (2-DG, glycolysis inhibitor, 50mM). X axis indicates measurement times. ECAR was normalized to 30,000 cells. data are presented as mean ± SE (n=3 for control; n=4 for IL-1β). **(B, C)** Representative graphs output from XF96 anlyzer showing the glycolysis (B) and glycolytic capacity (C). *p<0.05, and **p<0.01. **(D)** Effects of IL-1β on glucose uptake. 2-NBDG incorporation from organoids treated with PBS control (blue line) or IL-1β (red line) was determined by flow cytometry. Non-treated cells were used as a negative control for 2-NBDG treatment (grey-filled peak). **(E)** Representative histograms showing MFI (mean fluorescence of intensity) of 2-NBDG. Each individual dot represents one individual experiment and data are presented as mean ± SEM (n=4 for control; n=5 for IL-1β). ***p<0.001.

**Figure S8, related to Fig. 4H.**
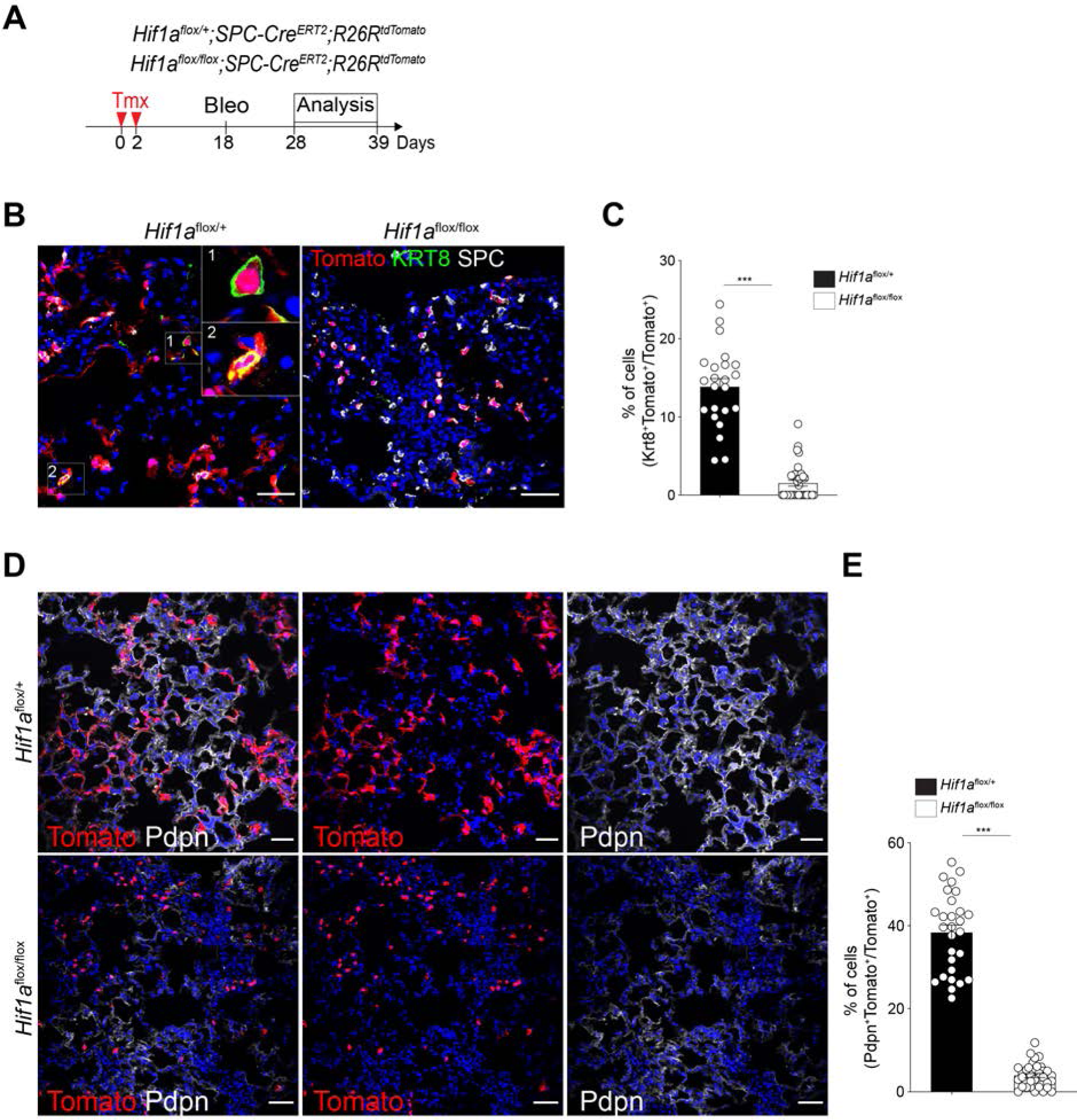
Deletion of *Hif1a* on AT2 cells impairs DATPs generation and AT1 cell regeneration. **(A)** Experimental design for lineage tracing. Date for analysis is as indicated. **(B)** Representative IF images showing *SPC* lineage-labeled DATPs at day 14 post injury in the lung of indicated genotyped mice: Tomato (for *SPC* lineage, red), Krt8 (green), SPC (white), and DAPI (blue). Insets (left) show high-power view (right top). Scale bars, 50μm. **(C)** Quantification of *SPC* lineage-labeled DATPs in (**B**). Each individual dot represents one section and data are presented as mean ± SEM with three independent experiments. Notably, there is a significant decrease in number of lineage-labeled DATPs in the absence of *Hif1a* in AT2 cells. **(D)** Representative IF images showing AT1 cell differentiation from *SPC* lineage-labeled cells at day 28 post injury in the lung of indicated genotyped mice: Tomato (for *SPC* lineage, red), Pdpn (white), and DAPI (blue). Scale bars, 50 μm. **(E)** Quantification of lineage-labeled Pdpn+ AT1 cells in (**D**). Each individual dot represents one section and data are presented as mean ± SEM (n=3 for Hif1a^flox/+^; n=4 for Hif1a^flox/flox^). Notably, there is a significant decrease in the number of lineage-labeled AT1 cells in the absence of *Hif1a* in AT2 cells.

**Figure S9, related to Fig. 5.**
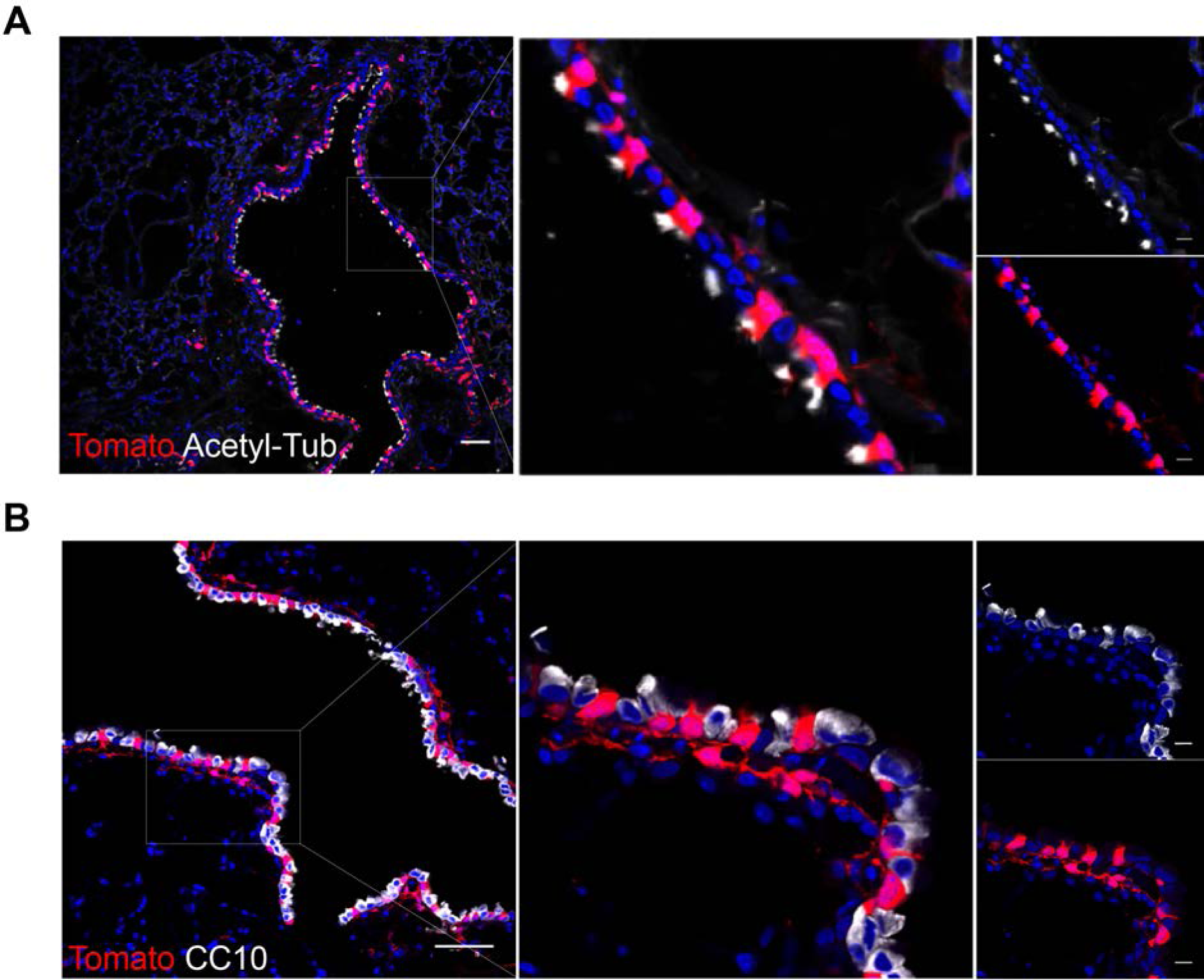
*Il1r1* lineage-labeled cells are found in airway ciliated cells in uninjured lungs. **(A, B)** Representative IF images showing *Il1r1* lineage-labeled cells only in ciliated cells (**A**) not in club cells (**B**) in uninjured lungs at day 14 post two doses of tamoxifen injection: Tomato (for *Il1r1* lineage, red), Acetyl-Tub (white), CC10 (white), and DAPI (blue). Scale bars: 50 μm.

**Figure S10, related to Fig. 6.**
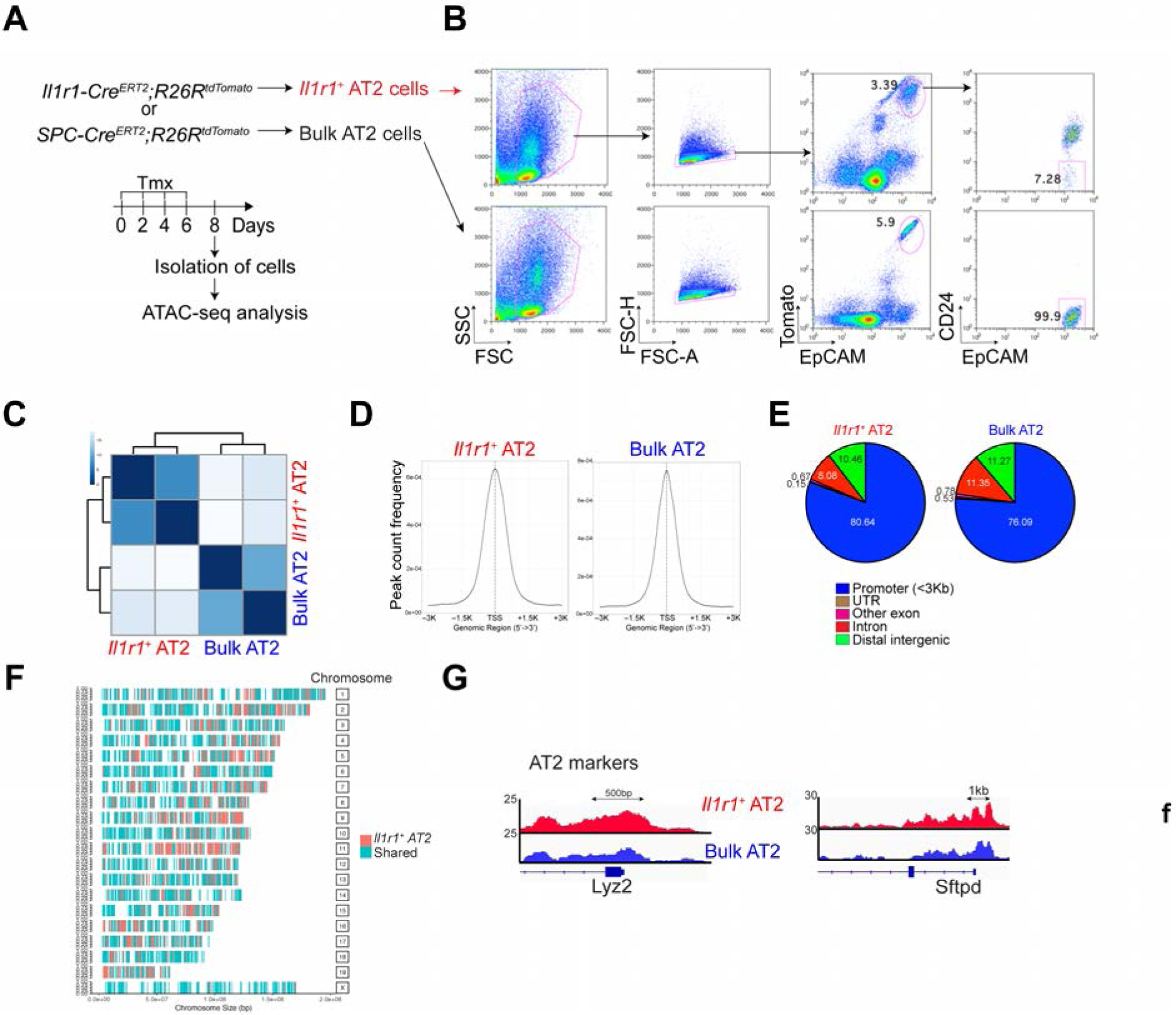
ATAC-seq analysis reveals distinct differences in open chromatin structure in *Il1r1*^+^AT2 cells versus bulk AT2 cells. **(A, B)** Experiment design (**A**) and sorting strategy by flow cytometry (**B**) for isolating *Il1r1*^+^AT2 or bulk AT2 cells from *Il1r1-Cre^ERT2^;R26R^tdTomato^* or *SPC-Cre^ERT2^;R26R^tdTomato^* mice, respectively. **(C)** Heat map of poisson distances between samples on the original count matrix. **(D)** Density plots depicting enrichment of ATAC-seq signals at TSSs ± 3 kb. **(E)** Distribution of ATAC-seq peaks within defined genomic regions of predicted mRNAs. UTR, untranslated regions. **(F)** Genome-wide profiling of ATAC-seq peaks in *Il1r1*^+^AT2 and bulk AT2 cells. **(G)** Snapshots of peaks enriched in shared genes Lyz2 and Sftpd. Arrows denote direction of transcription.

**Figure S11, related to Fig. 7, A-E.**
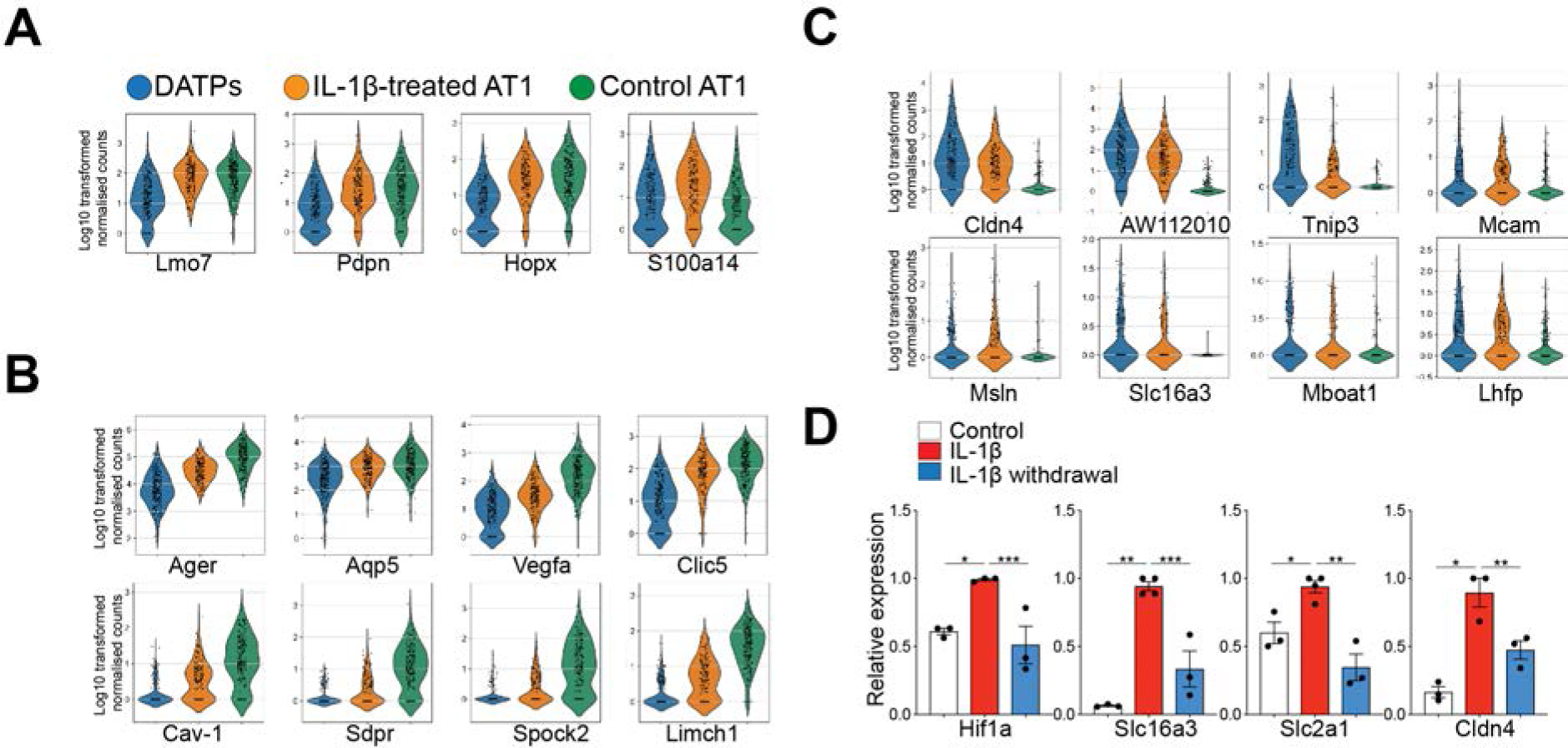
Sustained IL-1β prohibits mature AT1 differentiation by blocking the progression of DATPs into AT1 lineage. **(A-C)** Violin plots showing the log-transformed (log_10_(TPM+1)), normalized expression levels of early AT1 (**A**), late AT1 (**B**), and DATP (**C**) marker genes in DATPs, control or IL-1β-treated AT1 cells revealed by scRNA-seq analysis of organoids. **(D)** qPCR analysis of *Cldn4* and *Hif1a*-depenent genes that are involved in the glycolysis pathway after withdrawal of IL-1β in AT2 organoids. Each individual dot represents one experiment and date are presented as mean ± SEM. *p<0.5, **p<0.01, ***p<0.001.

**Figure S12, related to Fig. 7, F-H.**
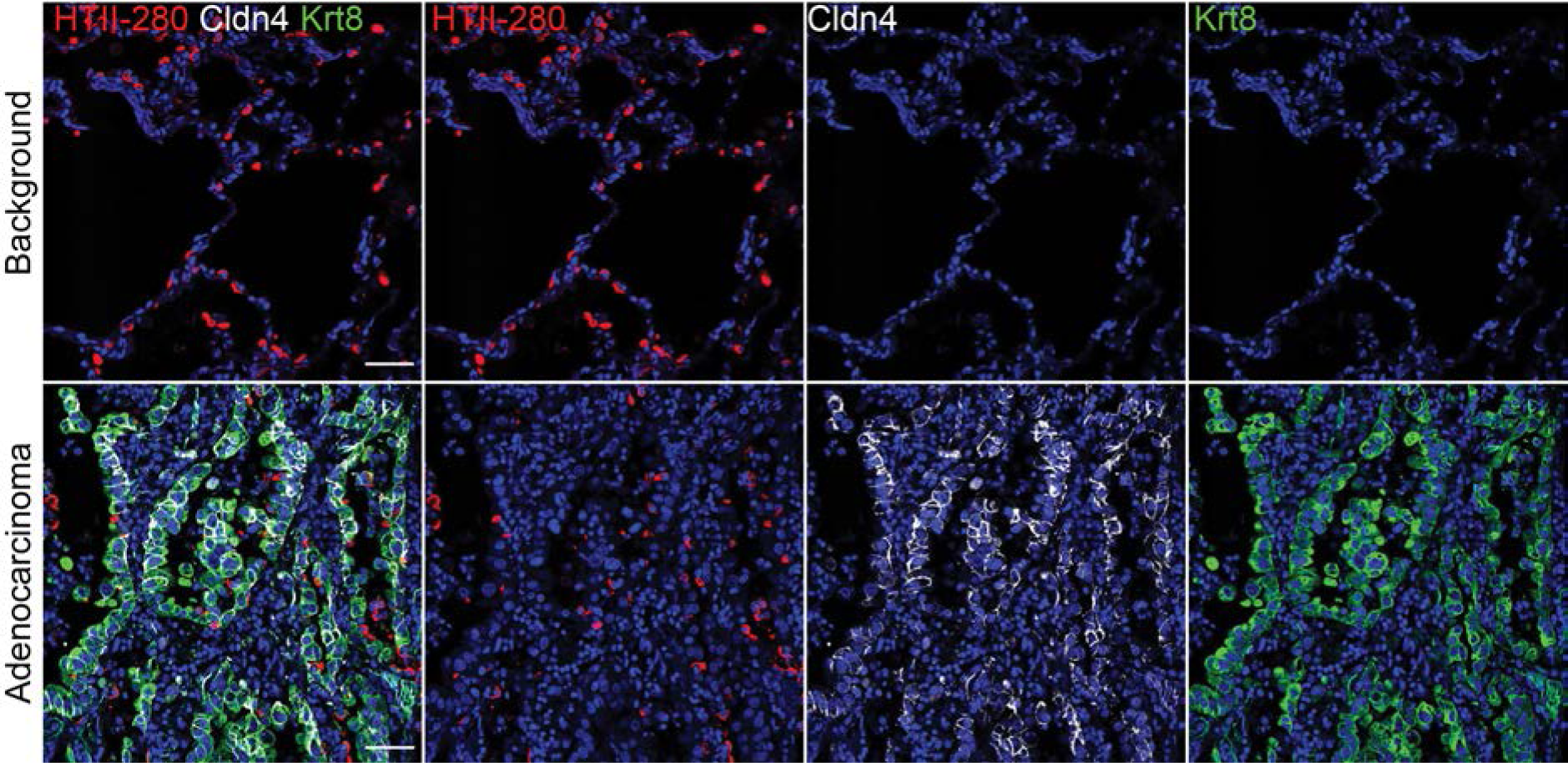
DATPs-like population is aberrantly accumulated in the lung from adenocarcinoma patients. Representative IF images of KRT8^+^CLDN4^+^ cells in the lung from adenocarcinoma patients (n=3). HTII-280 (red), CLDN4 (white), KRT8 (green) and DAPI (blue). Background region (top) in the lung tissue of the same patient was used for control. Scale bar, 50 μm.

## References

Adams, T.S., Schupp, J.C., Poli, S., Ayaub, E.A., Neumark, N., Ahangari, F., Chu, S.G., Raby, B.A., DeIuliis, G., Januszyk, M., et al. (2019). Single Cell RNA-seq reveals ectopic and aberrant lung resident cell populations in Idiopathic Pulmonary Fibrosis. bioRxiv https://doi.org/10.1101/759902.

Adamson, I.Y., and Bowden, D.H. (1974). The type 2 cell as progenitor of alveolar epithelial regeneration. A cytodynamic study in mice after exposure to oxygen. Lab Invest 30, 35–42.

Barkauskas, C.E., Cronce, M.J., Rackley, C.R., Bowie, E.J., Keene, D.R., Stripp, B.R., Randell, S.H., Noble, P.W., and Hogan, B.L. (2013). Type 2 alveolar cells are stem cells in adult lung. J Clin Invest 123, 3025–3036.

Bolger, A.M., Lohse, M., and Usadel, B. (2014). Trimmomatic: a flexible trimmer for Illumina sequence data. Bioinformatics 30, 2114–2120.

Buenrostro, J.D., Wu, B., Chang, H.Y., and Greenleaf, W.J. (2015). ATAC-seq: A Method for Assaying Chromatin Accessibility Genome-Wide. Curr Protoc Mol Biol 109, 21–29 21–29.

Chen, F., Liu, Z., Wu, W., Rozo, C., Bowdridge, S., Millman, A., Van Rooijen, N., Urban, J.F., Jr., Wynn, T.A., and Gause, W.C. (2012). An essential role for TH2-type responses in limiting acute tissue damage during experimental helminth infection. Nat Med 18, 260–266.

Conway, E.M., Pikor, L.A., Kung, S.H., Hamilton, M.J., Lam, S., Lam, W.L., and Bennewith, K.L. (2016). Macrophages, Inflammation, and Lung Cancer. Am J Respir Crit Care Med 193, 116–130.

Dang, C.V., Kim, J.W., Gao, P., and Yustein, J. (2008). The interplay between MYC and HIF in cancer. Nat Rev Cancer 8, 51–56.

Dobin, A., Davis, C.A., Schlesinger, F., Drenkow, J., Zaleski, C., Jha, S., Batut, P., Chaisson, M., and Gingeras, T.R. (2013). STAR: ultrafast universal RNA-seq aligner. Bioinformatics 29, 15–21.

Endo, T., Nakamura, J., Sato, Y., Asada, M., Yamada, R., Takase, M., Takaori, K., Oguchi, A., Iguchi, T., Higashi, A.Y., et al. (2015). Exploring the origin and limitations of kidney regeneration. J Pathol 236, 251–263.

Feng, J., Liu, T., Qin, B., Zhang, Y., and Liu, X.S. (2012). Identifying ChIP-seq enrichment using MACS. Nat Protoc 7, 1728–1740.

Finn, J., Sottoriva, K., Pajcini, K.V., Kitajewski, J.K., Chen, C., Zhang, W., Malik, A.B., and Liu, Y. (2019). Dlk1-Mediated Temporal Regulation of Notch Signaling Is Required for Differentiation of Alveolar TypeII to Type I Cells during Repair. Cell Rep 26, 2942–2954 e2945.

Fortier, M.A., Krishnaswamy, K., Danyod, G., Boucher-Kovalik, S., and Chapdalaine, P. (2008). A postgenomic integrated view of prostaglandins in reproduction: implications for other body systems. J Physiol Pharmacol 59 Suppl 1, 65–89.

Ganguly, K., Martin, T.M., Concel, V.J., Upadhyay, S., Bein, K., Brant, K.A., George, L., Mitra, A., Thimraj, T.A., Fabisiak, J.P., et al. (2014). Secreted phosphoprotein 1 is a determinant of lung function development in mice. Am J Respir Cell Mol Biol 51, 637–651.

Garayoa, M., Martinez, A., Lee, S., Pio, R., An, W.G., Neckers, L., Trepel, J., Montuenga, L.M., Ryan, H., Johnson, R., et al. (2000). Hypoxia-inducible factor-1 (HIF-1) up-regulates adrenomedullin expression in human tumor cell lines during oxygen deprivation: a possible promotion mechanism of carcinogenesis. Mol Endocrinol 14, 848–862.

Habermann, A.C., Gutierrez, A.J., Bui, L.T., Yahn, S.L., Winters, L.I., Calvi, C.L., Peter, L., Chung, M.I., Taylor, C.J., Jetter, C., et al. (2019). Single-cell RNA-sequencing reveals profibrotic roles of distinct epithelial and mesenchymal lineages in pulmonary fibrosis. bioRxiv https://doi.org/10.1101/753806.

Haghverdi, L., Buttner, M., Wolf, F.A., Buettner, F., and Theis, F.J. (2016). Diffusion pseudotime robustly reconstructs lineage branching. Nat Methods 13, 845–848.

Hasegawa, K., Sato, A., Tanimura, K., Uemasu, K., Hamakawa, Y., Fuseya, Y., Sato, S., Muro, S., and Hirai, T. (2017). Fraction of MHCII and EpCAM expression characterizes distal lung epithelial cells for alveolar type 2 cell isolation. Respir Res 18, 150.

Heinz, S., Benner, C., Spann, N., Bertolino, E., Lin, Y.C., Laslo, P., Cheng, J.X., Murre, C., Singh, H., and Glass, C.K. (2010). Simple combinations of lineage-determining transcription factors prime cis-regulatory elements required for macrophage and B cell identities. Mol Cell 38, 576–589.

Hogan, B.L., Barkauskas, C.E., Chapman, H.A., Epstein, J.A., Jain, R., Hsia, C.C., Niklason, L., Calle, E., Le, A., Randell, S.H., et al. (2014). Repair and regeneration of the respiratory system: complexity, plasticity, and mechanisms of lung stem cell function. Cell Stem Cell 15, 123–138.

Hsu, Y.C., Li, L., and Fuchs, E. (2014). Emerging interactions between skin stem cells and their niches. Nat Med 20, 847–856.

Katsura, H., Kobayashi, Y., Tata, P.R., and Hogan, B.L.M. (2019). IL-1 and TNFalpha Contribute to the Inflammatory Niche to Enhance Alveolar Regeneration. Stem Cell Reports 12, 657–666.

Klose, C.S., and Artis, D. (2016). Innate lymphoid cells as regulators of immunity, inflammation and tissue homeostasis. Nat Immunol 17, 765–774.

Kobayashi, Y., Tata, A., Konkimalla, A., Katsura, H., Lee, F.R., Ou, J., Banovich, E.N., Kropski, A.J., and Tata, R.P. (2019). Persistence of a novel regeneration-associated transitional cell state in pulmonary fibrosis. BioRxiv https://doi.org/10.1101/855155.

Kotton, D.N., and Morrisey, E.E. (2014). Lung regeneration: mechanisms, applications and emerging stem cell populations. Nat Med 20, 822–832.

Kuriakose, T., and Kanneganti, T.D. (2018). ZBP1: Innate Sensor Regulating Cell Death and Inflammation. Trends Immunol 39, 123–134.

Lechner, A.J., Driver, I.H., Lee, J., Conroy, C.M., Nagle, A., Locksley, R.M., and Rock, J.R. (2017). Recruited Monocytes and Type 2 Immunity Promote Lung Regeneration following Pneumonectomy. Cell Stem Cell 21, 120–134 e127.

Lee, J.H., Bhang, D.H., Beede, A., Huang, T.L., Stripp, B.R., Bloch, K.D., Wagers, A.J., Tseng, Y.H., Ryeom, S., and Kim, C.F. (2014). Lung stem cell differentiation in mice directed by endothelial cells via a BMP4-NFATc1-thrombospondin-1 axis. Cell 156, 440–455.

Lee, J.H., Tammela, T., Hofree, M., Choi, J., Marjanovic, N.D., Han, S., Canner, D., Wu, K., Paschini, M., Bhang, D.H., et al. (2017). Anatomically and Functionally Distinct Lung Mesenchymal Populations Marked by Lgr5 and Lgr6. Cell 170, 1149–1163 e1112.

Li, L., and Clevers, H. (2010). Coexistence of quiescent and active adult stem cells in mammals. Science 327, 542–545.

Ligresti, G., Aplin, A.C., Dunn, B.E., Morishita, A., and Nicosia, R.F. (2012). The acute phase reactant orosomucoid-1 is a bimodal regulator of angiogenesis with time- and context-dependent inhibitory and stimulatory properties. PLoS One 7, e41387.

Lindemans, C.A., Calafiore, M., Mertelsmann, A.M., O'Connor, M.H., Dudakov, J.A., Jenq, R.R., Velardi, E., Young, L.F., Smith, O.M., Lawrence, G., et al. (2015). Interleukin-22 promotes intestinal-stem-cell-mediated epithelial regeneration. Nature 528, 560–564.

Madisen, L., Zwingman, T.A., Sunkin, S.M., Oh, S.W., Zariwala, H.A., Gu, H., Ng, L.L., Palmiter, R.D., Hawrylycz, M.J., Jones, A.R., et al. (2010). A robust and high-throughput Cre reporting and characterization system for the whole mouse brain. Nat Neurosci 13, 133–140.

Mantovani, A., Allavena, P., Sica, A., and Balkwill, F. (2008). Cancer-related inflammation. Nature 454, 436–444.

Martis, P.C., Whitsett, J.A., Xu, Y., Perl, A.K., Wan, H., and Ikegami, M. (2006). C/EBPalpha is required for lung maturation at birth. Development 133, 1155–1164.

Maynard, A., McCoach, C.E., Julia K. Rotow, L.H., Haderk, F., Kerr, L., Yu, E.A., Schenk, E.L., Tan, W., Zee, A., Tan, M., et al. (2019). Heterogeneity and targeted therapy-induced adaptations in lung cancer revealed by longitudinal single-cell RNA sequencing. bioRxiv doi: https://doi.org/10.1101/2019.12.08.868828.

Medzhitov, R. (2008). Origin and physiological roles of inflammation. Nature 454, 428–435.

Minoo, P., Su, G., Drum, H., Bringas, P., and Kimura, S. (1999). Defects in tracheoesophageal and lung morphogenesis in Nkx2.1(-/-) mouse embryos. Dev Biol 209, 60–71.

Miossec, P., and Kolls, J.K. (2012). Targeting IL-17 and TH17 cells in chronic inflammation. Nat Rev Drug Discov 11, 763–776.

Misharin, A.V., Morales-Nebreda, L., Reyfman, P.A., Cuda, C.M., Walter, J.M., McQuattie-Pimentel, A.C., Chen, C.I., Anekalla, K.R., Joshi, N., Williams, K.J.N., et al. (2017). Monocyte-derived alveolar macrophages drive lung fibrosis and persist in the lung over the life span. J Exp Med 214, 2387–2404.

Moll, H.P., Pranz, K., Musteanu, M., Grabner, B., Hruschka, N., Mohrherr, J., Aigner, P., Stiedl, P., Brcic, L., Laszlo, V., et al. (2018). Afatinib restrains K-RAS-driven lung tumorigenesis. Sci Transl Med 10.

Nabhan, A.N., Brownfield, D.G., Harbury, P.B., Krasnow, M.A., and Desai, T.J. (2018). Single-cell Wnt signaling niches maintain stemness of alveolar type 2 cells. Science 359, 1118–1123.

Naik, S., Larsen, S.B., Cowley, C.J., and Fuchs, E. (2018). Two to Tango: Dialog between Immunity and Stem Cells in Health and Disease. Cell 175, 908–920.

Naik, S., Larsen, S.B., Gomez, N.C., Alaverdyan, K., Sendoel, A., Yuan, S., Polak, L., Kulukian, A., Chai, S., and Fuchs, E. (2017). Inflammatory memory sensitizes skin epithelial stem cells to tissue damage. Nature 550, 475–480.

Olsson, A., Venkatasubramanian, M., Chaudhri, V.K., Aronow, B.J., Salomonis, N., Singh, H., and Grimes, H.L. (2016). Single-cell analysis of mixed-lineage states leading to a binary cell fate choice. Nature 537, 698–702.

Ramirez, F., Ryan, D.P., Gruning, B., Bhardwaj, V., Kilpert, F., Richter, A.S., Heyne, S., Dundar, F., and Manke, T. (2016). deepTools2: a next generation web server for deep-sequencing data analysis. Nucleic Acids Res 44, W160–165.

Riemondy, K.A., Jansing, N.L., Jiang, P., Redente, E.F., Gillen, A.E., Fu, R., Miller, A.J., Spence, J.R., Gerber, A.N., Hesselberth, J.R., et al. (2019). Single cell RNA sequencing identifies TGFbeta as a key regenerative cue following LPS-induced lung injury. JCI Insight 5.

Rindler, T.N., Stockman, C.A., Filuta, A.L., Brown, K.M., Snowball, J.M., Zhou, W., Veldhuizen, R., Zink, E.M., Dautel, S.E., Clair, G., et al. (2017). Alveolar injury and regeneration following deletion of ABCA3. JCI Insight 2.

Robson, M.J., Zhu, C.B., Quinlan, M.A., Botschner, D.A., Baganz, N.L., Lindler, K.M., Thome, J.G., Hewlett, W.A., and Blakely, R.D. (2016). Generation and Characterization of Mice Expressing a Conditional Allele of the Interleukin-1 Receptor Type 1. PLoS One 11, e0150068.

Rock, J.R., Barkauskas, C.E., Cronce, M.J., Xue, Y., Harris, J.R., Liang, J., Noble, P.W., and Hogan, B.L. (2011). Multiple stromal populations contribute to pulmonary fibrosis without evidence for epithelial to mesenchymal transition. Proc Natl Acad Sci U S A 108, E1475–1483.

Schonthaler, H.B., Guinea-Viniegra, J., and Wagner, E.F. (2011). Targeting inflammation by modulating the Jun/AP-1 pathway. Ann Rheum Dis 70 Suppl 1, i109–112.

Semenza, G.L. (2012). Hypoxia-inducible factors in physiology and medicine. Cell 148, 399–408.

Van Keymeulen, A., Rocha, A.S., Ousset, M., Beck, B., Bouvencourt, G., Rock, J., Sharma, N., Dekoninck, S., and Blanpain, C. (2011). Distinct stem cells contribute to mammary gland development and maintenance. Nature 479, 189–193.

Wagers, A.J., and Weissman, I.L. (2004). Plasticity of adult stem cells. Cell 116, 639–648.

Weaver, M., Dunn, N.R., and Hogan, B.L. (2000). Bmp4 and Fgf10 play opposing roles during lung bud morphogenesis. Development 127, 2695–2704.

Westphalen, K., Gusarova, G.A., Islam, M.N., Subramanian, M., Cohen, T.S., Prince, A.S., and Bhattacharya, J. (2014). Sessile alveolar macrophages communicate with alveolar epithelium to modulate immunity. Nature 506, 503–506.

Wolf, F.A., Angerer, P., and Theis, F.J. (2018). SCANPY: large-scale single-cell gene expression data analysis. Genome Biol 19, 15.

Wolf, F.A., Hamey, F.K., Plass, M., Solana, J., Dahlin, J.S., Gottgens, B., Rajewsky, N., Simon, L., and Theis, F.J. (2019). PAGA: graph abstraction reconciles clustering with trajectory inference through a topology preserving map of single cells. Genome Biol 20, 59.

Wu, H., Yu, Y., Huang, H., Hu, Y., Fu, S., Wang, Z., Shi, M., Zhao, X., Yuan, J., Li, J., et al. (2020). Progressive Pulmonary Fibrosis Is Caused by Elevated Mechanical Tension on Alveolar Stem Cells. Cell 180, 107–121 e117.

Yu, G., Wang, L.G., and He, Q.Y. (2015). ChIPseeker: an R/Bioconductor package for ChIP peak annotation, comparison and visualization. Bioinformatics 31, 2382–2383.

Zacharias, W.J., Frank, D.B., Zepp, J.A., Morley, M.P., Alkhaleel, F.A., Kong, J., Zhou, S., Cantu, E., and Morrisey, E.E. (2018). Regeneration of the lung alveolus by an evolutionarily conserved epithelial progenitor. Nature 555, 251–255.

Zeng, L., Yang, X.T., Li, H.S., Li, Y., Yang, C., Gu, W., Zhou, Y.H., Du, J., Wang, H.Y.,Sun, J.H., et al. (2016). The cellular kinetics of lung alveolar epithelial cells and its relationship with lung tissue repair after acute lung injury. Respir Res 17, 164.

Zepp, J.A., Zacharias, W.J., Frank, D.B., Cavanaugh, C.A., Zhou, S., Morley, M.P., and Morrisey, E.E. (2017). Distinct Mesenchymal Lineages and Niches Promote Epithelial Self-Renewal and Myofibrogenesis in the Lung. Cell 170, 1134–1148 e1110.

Zhang, Z., Newton, K., Kummerfeld, S.K., Webster, J., Kirkpatrick, D.S., Phu, L., Eastham-Anderson, J., Liu, J., Lee, W.P., Wu, J., et al. (2017). Transcription factor Etv5 is essential for the maintenance of alveolar type II cells. Proc Natl Acad Sci U S A 114, 3903–3908.

